# A paralog of a clonal propagation regulator promotes cell-cycle re-entry during thallus regeneration in *Marchantia polymorpha*

**DOI:** 10.64898/2026.08.26.747441

**Authors:** Yukiko Yasui, Hirotaka Kato, Yuuki Sakai, Gaku Konishi, Suzuha Tanaka, Hidehiro Fukaki, Tetsuro Mimura, Ryuichi Nishihama, Takayuki Kohchi, Kimitsune Ishizaki

## Abstract

Plants possess a remarkable capacity for regeneration, which involves the redeployment of developmental programs and diverse regulatory mechanisms. However, how related regulators with overlapping functions are differentially deployed during regeneration remains poorly understood. The model liverwort *Marchantia polymorpha* provides a powerful experimental system for studying regeneration because it readily regenerates apical meristems from basal thallus fragments after removal of the original meristem, even without exogenous plant hormones. Here, we identify the R2R3-MYB transcription factor GEMMA CUP-ASSOCIATED MYB1-LIKE (MpGC1L), the closest paralog of the clonal propagation regulator MpGCAM1, as a positive regulator of regeneration. MpGC1L was rapidly induced at the cut site following meristem removal. Ectopic overexpression of MpGC1L caused the proliferation of undifferentiated cells, whereas *Mpgc1l* mutants showed delayed regeneration and reduced S-phase entry. Loss of MpGCAM1 alone had little effect on regeneration but markedly enhanced the *Mpgc1l* phenotype, indicating partially redundant functions. Transcriptome analysis of the double mutant revealed reduced induction of genes associated with ribosome biogenesis and the cell cycle. We next examined the relationship between MpGC1L and the known jasmonate- and auxin- related regeneration regulators, MpERF15 and MpLAXR. Mp*GC1L* induction was retained in Mp*erf15* and Mp*laxr* mutants and was unaffected by OPDA or auxin treatment, whereas Mp*ERF15* and Mp*LAXR* were still induced in Mp*gc1l* Mp*gcam1* double mutants. Thus, these regulators are not arranged in a simple linear transcriptional pathway. Our findings reveal that the paralogous MYB transcription factors MpGC1L and MpGCAM1 promote cell proliferation in distinct developmental contexts, thereby linking clonal propagation and wound-induced regeneration.

## Introduction

Plants rely on adaptive strategies distinct from those of animals. Plants cannot move as animals do to adjust to their environments. Instead, plants deploy indeterminate development; they maintain stem cells within meristems, allowing them to continuously develop organs throughout their lifecycles ^1,2^. In addition, plants can initiate new meristems, such as secondary meristems, during normal development ^3^ or reestablish meristems *de novo* during regeneration ^4,5^. These remarkable abilities are vital for plant survival.

To date, research on plant regeneration has primarily focused on the model angiosperm *Arabidopsis thaliana*. *A. thaliana* exhibits various forms of regeneration, including the *de novo* formation of the shoot apical meristem and root apical meristem from differentiated tissues, which give rise to the aerial and underground parts of the plant body, respectively. Numerous studies have investigated the underlying molecular mechanisms, particularly the reactivation of normal developmental programs. For example, transcription factors involved in lateral root formation, such as PLETHORA (PLT), a member of the APETALA2/ETHYLENE RESPONSE FACTOR (AP2/ERF) transcription factor family, play crucial roles in callus formation, a mass of dedifferentiated cells, and the acquisition of competence for shoot regeneration ^6,7^. Additionally, the ectopically induced expression of genes encoding shoot meristem regulators, such as ENHANCER OF SHOOT REGENERATION 1 (ESR1), another AP2/ERF transcription factor, in root explants leads to *de novo* shoot regeneration ^8–10^.

Several phytohormones also play important roles in plant regeneration. Treatment with auxin and cytokinin enhances the regenerative capacity of *A. thaliana* ^11^. For efficient shoot regeneration, callus formation is first induced by incubating root or hypocotyl explants in auxin-rich medium. The calli are then transferred to cytokinin-rich medium to promote adventitious shoot formation. Plants can initiate a regeneration process in response to injury ^4^. Recent studies have revealed that jasmonate, a well-characterized wounding-associated plant hormone recognized by the F-box protein CORONATINE INSENSITIVE1 (COI1), links wounding to specific types of regeneration. This process involves AP2/ERF transcription factors such as ERF109 and ERF115, which mediate regeneration following wounding ^12–14^. Thus, many AP2/ERF transcription factors involved in regeneration have been identified in *A. thaliana*.

Bryophytes, including liverworts, which are the sister to vascular plants, exhibit a haploid-dominant lifecycle and are known for their high regenerative capacity ^15^. During the haploid gametophyte generation in the model liverwort *Marchantia polymorpha*, the plant body grows via an apical meristem located in the notch at the thallus tip ^16^. If the apical meristem is removed, the basal explant regenerates new apical meristems, primarily from its apical end, even in the absence of plant hormone treatment ^17,18^.

LOW-AUXIN RESPONSIVE (MpLAXR), an AP2/ERF transcription factor and a putative ortholog of ESR1, was recently identified as a key regulator of cellular reprogramming during regeneration in *M. polymorpha* ^19,20^. In intact plant bodies, Mp*LAXR* is downregulated by auxin. Upon removal of the apical meristem, auxin levels decrease, leading to the upregulation of Mp*LAXR* in the basal region. This upregulation subsequently triggers cellular reprogramming ^19^.

Another AP2/ERF transcription factor gene, Mp*ERF15*, an ortholog of *A. thaliana ERF109*, is rapidly upregulated in response to wounding ^21^. MpERF15 contributes to the biosynthesis of 12-oxo-phytodienoic acid (OPDA) and dinor-OPDA (dn-OPDA). dn-iso-OPDA and Δ 4-dn-iso-OPDA act as ligands for MpCOI1, whereas OPDA serves as a biosynthetic precursor of dn-OPDA in *M. polymorpha* ^22–24^. MpERF15 promotes the expression of genes encoding 13-lipoxygenase (LOX), allene oxide synthase (AOS), and allene oxide cyclase (AOC) ^21,22^. OPDA and dn-OPDA positively regulate Mp*ERF15* expression via an MpCOI1-dependent signaling pathway, suggesting the presence of a positive feedback loop between MpERF15 and the biosynthesis of OPDA and dn-OPDA ^21^. MpERF15 also positively regulates Mp*LAXR* expression during regeneration ^21^. Despite the critical roles of MpLAXR and MpERF15 in regeneration, mutants of both genes exhibit delayed regeneration rather than a complete loss of regenerative ability ^19,21^. This suggests that the molecular mechanisms underlying efficient regeneration in *M. polymorpha* have yet to be fully elucidated.

In addition to its remarkable capacity for regeneration, *M. polymorpha* exhibits a strong ability to generate new stem cells during normal development. During the vegetative stage of the gametophyte generation, the plant body (thallus) undergoes periodic bifurcation, in which single apical meristem gives rise to two. ^16,25^. Also, on the dorsal side of the thallus, specialized organs for vegetative reproduction known as gemma cups form. Within each gemma cup, more than 100 clonal propagules (gemmae) are produced, each containing two meristems.

The R2R3-MYB transcription factor GEMMA CUP-ASSOCIATED MYB1 (MpGCAM1) is an essential regulator of gemma and gemma cup formation. MpGCAM1 promotes the proliferation of undifferentiated cells, thereby facilitating the development of these structures ^26,27^. In this study, we identified MpGC1L, a closest paralog of the gemma cup regulator MpGCAM1, as a novel factor involved in regeneration in *M. polymorpha*. MpGC1L expression was rapidly induced in basal explants following apical meristem removal, and its overexpression promoted the proliferation of undifferentiated cells. While loss-of-function mutants of MpGC1L showed delayed regeneration, double mutants lacking both MpGC1L and MpGCAM1 exhibited severe defects. Notably, MpGC1L and MpGCAM1 function independently of known regeneration regulators MpLAXR and MpERF15, suggesting the existence of a parallel regeneration pathway in *M. polymorpha*.

## Results

### Mp*GC1L* is upregulated at the regeneration site

Mp*GCAM1*, a key contributor to the proliferation of undifferentiated cells during gemma cup and gemma formation, belongs to R2R3-MYB subfamily 14 ^26^. This subfamily also includes *REGULATOR OF AXILLARY MERISTEMs* (*RAXs*), which regulate axillary meristem formation in *A. thaliana* ^28,29^. In *M. polymorpha,* another member of this subfamily, Mp*R2R3-MYB20* (Mp4g21790), has been identified ^26^ and named Mp*GC1L*.

To investigate the functions of Mp*GC1L*, we first examined its tissue-specific expression pattern using the *M. polymorpha* MarpolBase Expression database (MBEX) ^30^. Unlike Mp*GCAM1*, Mp*GC1L* was expressed at low levels in gemma cups but at higher levels in basal thallus fragments 24 hours after removing the apical meristem ^26,31^ (Figure S1). We measured Mp*GC1L* expression at the regeneration site using RT-qPCR in a time-course experiment. Mp*GC1L* expression sharply increased in basal explants 1 h after meristem removal and remained consistently high for at least 15 hours in wild-type (WT) plants (Figure 1A).

**Figure 1.**
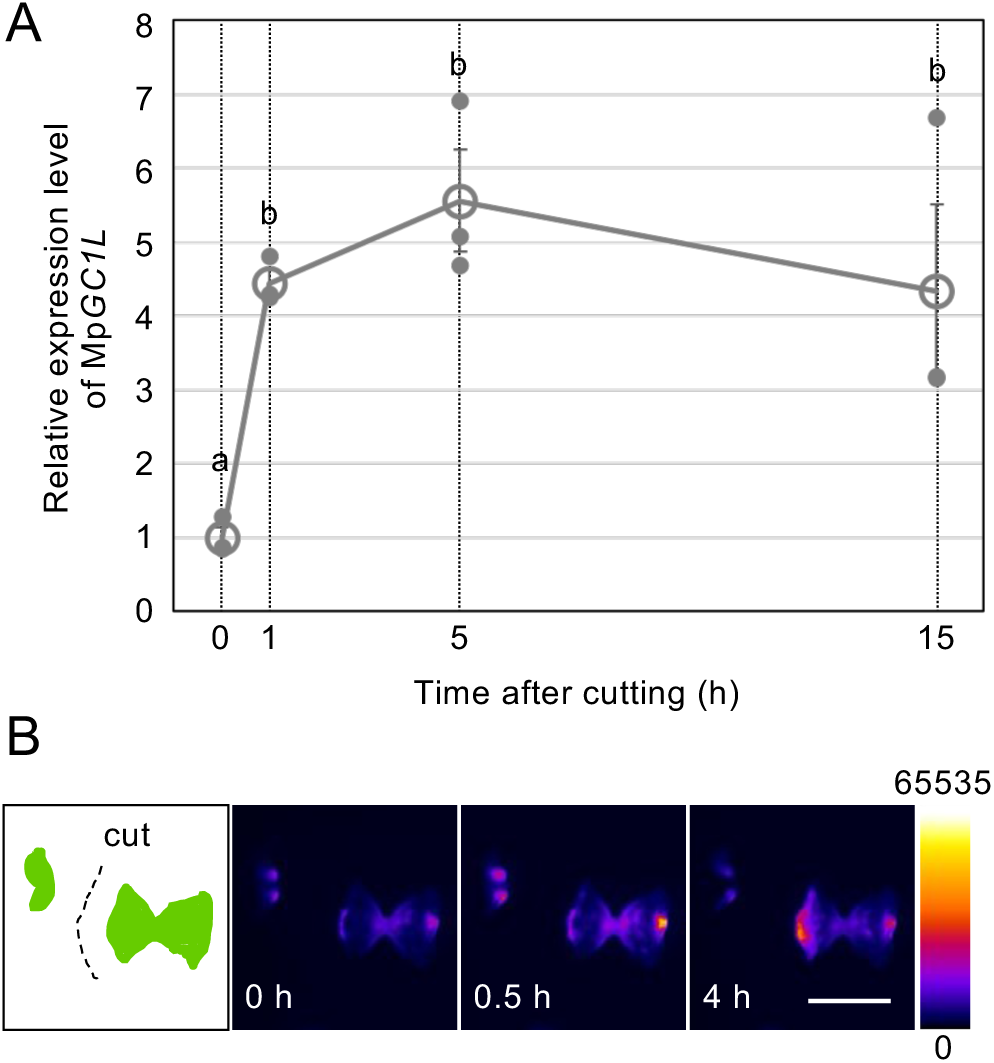
Mp*GC1L* expression levels during regeneration. (A) RT-qPCR analysis of Mp*GC1L* expression in basal explants after cutting off the apical meristem. 10-d-old wild-type (WT) thalli were cut, and basal explants were cultured for the indicated time. Mp*EF1α* was used for normalization. Data are mean ± SE. Different letters above the markers indicate significant differences at *p* < 0.05 by Tukey–Kramer test (*n* = 3 biological replicates). (B) Promoter activity of Mp*GC1L* during thallus regeneration. One apical side of a 10-day-old Mp*GC1Lpro:ELUC* gemmaling was removed (left), and luciferase activity was measured at 0, 0.5, or 4 h. Relative luciferase activity is pseudo-colored as shown in the right panel. Scale bar = 5 mm.

To further examine the temporal and spatial expression pattern of Mp*GC1L* at the regeneration site, we generated transgenic plants expressing luciferase under the control of the Mp*GC1L* promoter. Luminescence signals began to increase within 30 minutes at the cut site of the basal explant following meristem excision and peaked approximately 4 hours later (Figure 1B), which is consistent with the results of RT-qPCR. These findings suggest that MpGC1L plays a role in regeneration in *M. polymorpha*.

### MpGC1L promotes the proliferation of undifferentiated cells

Given that MpGCAM1 is known to promote the proliferation of undifferentiated cells ^26^, we reasoned that MpGC1L might have similar functions. To test this hypothesis, we generated transgenic *M. polymorpha* plants overexpressing MpGC1L fused to the N-terminus of a glucocorticoid receptor (GR) domain under the control of the Mp*ELONGATION FACTOR 1α* promoter (Mp*EFpro:*Mp*GC1L-GR*). In these transgenic plants, MpGC1L function could be induced by dexamethasone (DEX) treatment. Mock-treated Mp*EFpro:*Mp*GC1L-GR* plants exhibited no abnormalities and showed a WT-like phenotype. The plants developed air chambers and gemma cups on the dorsal side, while rhizoids were initiated on the ventral side at the apical notch (Figures 2 and S2) ^16^. By contrast, DEX-treated Mp*EFpro:*Mp*GC1L-GR* plants displayed an undifferentiated cell mass phenotype, forming numerous irregular structures and only a few rhizoids while failing to develop apical notches or normal dorsal/ventral axes (Figures 2 and S2). These findings suggest that MpGC1L promotes the proliferation of undifferentiated cells while simultaneously inhibiting cell differentiation.

**Figure 2.**
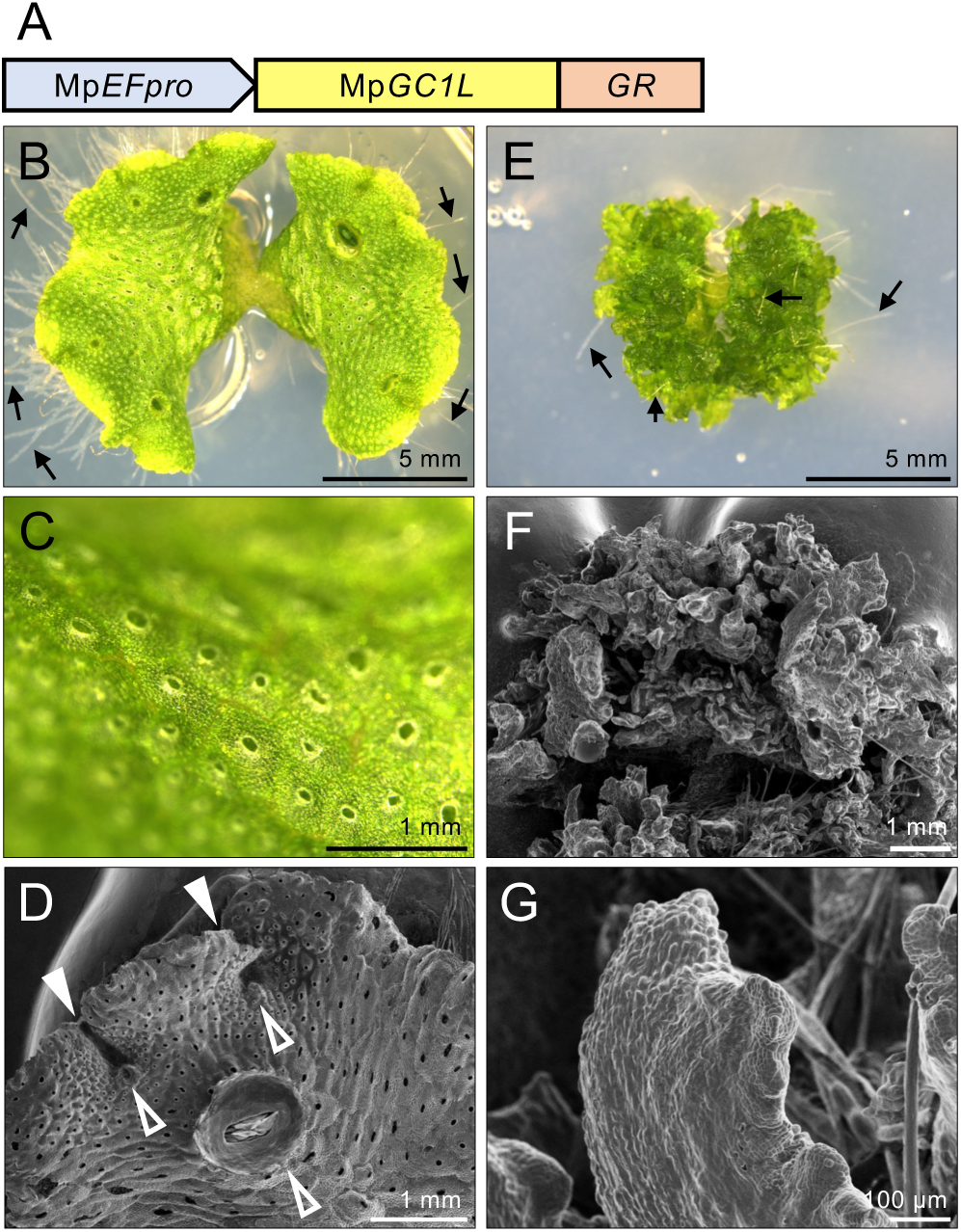
Induction of the ectopic overexpression of MpGC1L function. (A) Schematic representation of the Mp*EFpro:*Mp*GC1L-GR* construct. (B-D) Mock-treated 2-week-old Mp*EFpro:*Mp*GC1L-GR* #4 transgenic plants. Image of a whole plant (B). The thallus surface showing air chambers (C). Scanning electron microscopy image (D). (E-G) Two-week-old Mp*EFpro:*Mp*GC1L-GR* #4 plants treated with 10 μM DEX. Whole plant (E), and scanning electron microscopy images (F and G). Arrows in (B) and (E) indicate rhizoids. Filled and open arrowheads in (D) indicate apical notches and gemma cups, respectively.

### MpGC1L plays a pivotal role in regeneration, with a subsidiary function of MpGCAM1

To further investigate the roles of MpGC1L in plant regeneration, we generated Mp*gc1l* mutants using CRISPR/Cas9-mediated targeted genome editing. Two independent Mp*gc1l^ge^* mutant lines were established, each targeting different sequences. Neither mutant exhibited obvious morphological abnormalities, including gemma cup formation, indicating that MpGC1L is not involved in gemma cup development, unlike MpGCAM1, although one mutant showed a slightly reduced growth rate compared with the WT at later stages of growth (Figure S3).

Given that Mp*GC1L* is upregulated at the regeneration site in WT plants (Figure 1), we analyzed the regeneration of these Mp*gc1l^ge^*mutants. After gemmae were grown, we excised their apical meristems, cultured the basal explants, and observed regeneration in these explants. In WT plants, unequivocal regenerative buds formed 7 days after excision. However, in the Mp*gc1l^ge^* mutants, only poorly defined buds were observed at this time point (Figures 3A-3C and 3G). By 10 days after excision, regenerative buds had emerged in the mutants, but they were significantly smaller than those of WT plants (Figures 3D-3F and 3H). These findings indicate that regeneration is significantly delayed in the Mp*gc1l^ge^* mutants.

**Figure 3.**
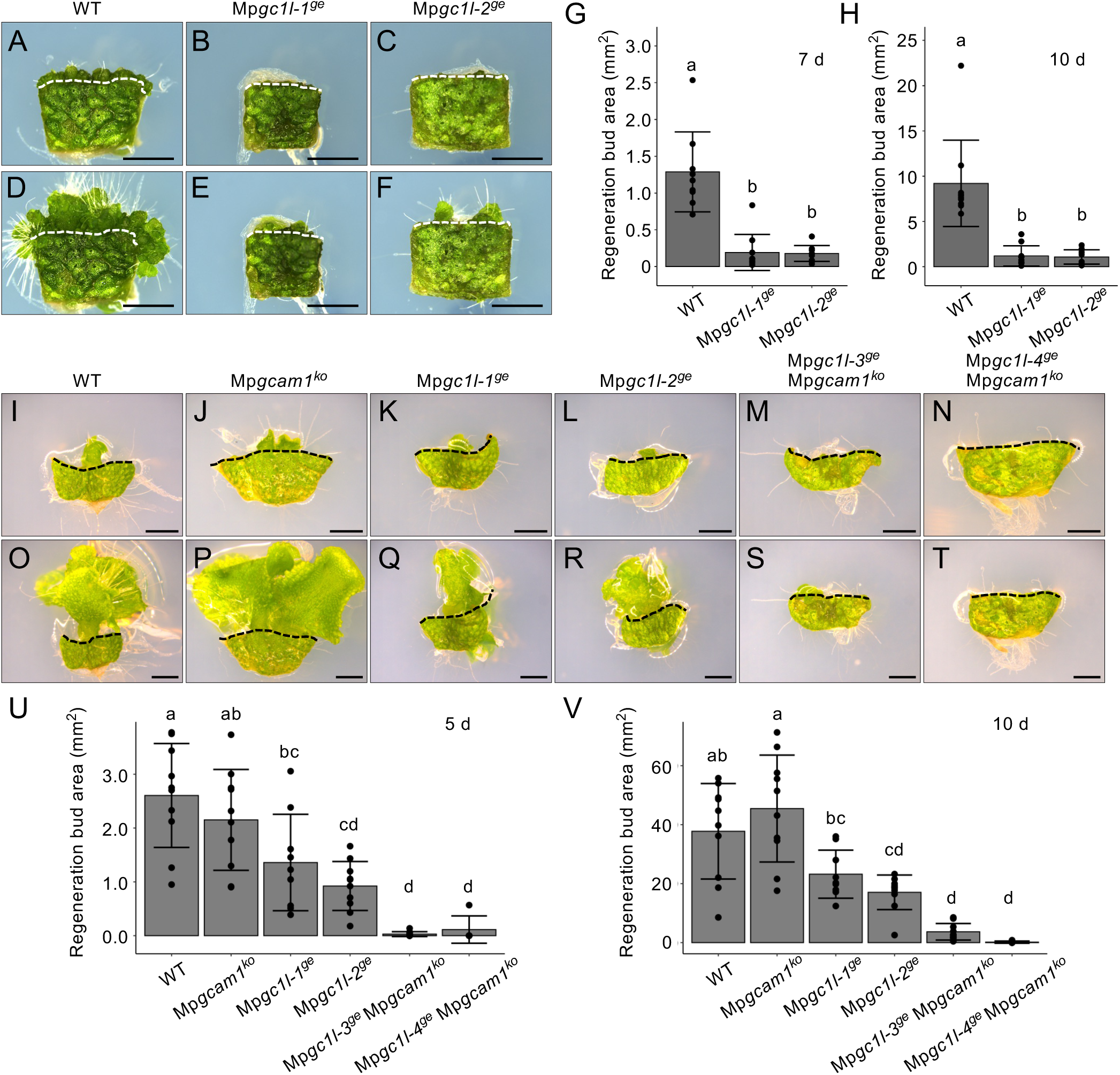
Regeneration of Mp*gcam1,* Mp*gc1l*, and double mutants. (A-F) Observation of regenerative bud formation in Mp*gc1l* single mutant gemmalings 7 d (A-C) and 10 d (D-F) after cutting off the meristem. Bars = 2 mm. WT (A and D), Mp*gc1l-1^ge^* (B and E), Mp*gc1l-2^ge^* (C and F). The dotted lines show the border between the basal explant and regenerated region. (G and H) Quantification of regenerative bud area using gemmalings 7 d (G) and 10 d (H) after cutting off the apical meristem. Different letters above the bars indicate significant differences at *p* < 0.01 by Tukey–Kramer test. n = 10. (I-T) Observation of regenerative bud formation in thalli 5 d (I-N) and 10 d (O-T) after cutting off the apical meristem. Bars = 2 mm. WT (I and O), Mp*gcam1^ko^* (J and P), Mp*gc1l-1^ge^* (K and Q), Mp*gc1l-2^ge^* (L and R), Mp*gc1l-3^ge^ gcam1^ko^* (M and S), and Mp*gc1l-4^ge^ gcam1^ko^* (N and T). The dotted lines show the border between the basal explant and regenerated region. (U and V) Quantification of regenerative bud area using thalli 5 d (U) and 10 d (V) after cutting off the apical meristem. Different letters above the bars indicate significant differences *p* < 0.05 by Tukey–Kramer test. *n* = 10 (WT, Mp*gcam1^ko^*, Mp*gc1l-1^ge^*, Mp*gc1l-2^ge^*, Mp*gc1l-3^ge^* Mp*gcam1^ko^*); *n* = 5 (Mp*gc1l-4^ge^* Mp*gcam1^ko^*).

Due to sequence similarity of the R2R3-MYB domains between MpGC1L and MpGCAM1 (Figure S4), we reasoned that MpGCAM1 might also contribute to regeneration. To test this hypothesis, we disrupted Mp*GC1L* in the Mp*gcam1^ko^* mutant background ^26^ by CRISPR/Cas9-mediated genome editing using the same target sequences used to generate the single Mp*gc1l^ge^*mutants (Figure S4). Since the Mp*gcam1^ko^* mutant did not produce gemmae, we grew plants from the tips of thalli and excised their apical meristems to generate basal explants. Under these conditions, the Mp*gc1l^ge^*basal explants developed significantly smaller regenerative buds compared to WT (Figures 3I-3V), although the difference was less pronounced than that using gemmalings (Figures 3A-3H). Regeneration in Mp*gcam1^ko^* was similar to that of the WT. By contrast, the Mp*gc1l^ge^* Mp*gcam1^ko^* double mutants showed severely delayed regeneration compared to both WT and Mp*gc1l^ge^*single mutant plants (Figures 3I-3V). Furthermore, in apical growth, meristem activity was occasionally exhausted in the double mutants (Figure S4).

Next, we measured the expression levels of Mp*GCAM1* during regeneration in both WT and Mp*gc1l^ge^* plants by RT-qPCR. Mp*GCAM1* expression did not change significantly 1 hour after excision but increased at later time points (Figure S5). These findings indicate that MpGC1L makes the predominant contribution to regeneration under these conditions, whereas MpGCAM1 provides partially redundant activity that becomes evident in the absence of MpGC1L.

### The pathway downstream of MpGC1L is essential for regeneration

To investigate the effects of repressing genes downstream of MpGC1L, we used the SUPERMAN repressor domain X (SRDX) system ^32^. We generated Mp*EFpro:*Mp*GC1L-SRDX-GR* transgenic plants in the WT background, in which MpGC1L-SRDX functions as a dominant repressor that can be ectopically induced by DEX treatment (Figure S6). In DEX-treated Mp*EFpro:*Mp*GC1L-SRDX-GR* plants, overall plant growth was inhibited compared to mock-treated plants. However, tissue differentiation, such as rhizoid and air chamber formation, was still observed (Figures S6B-S6E). In mock-treated plants, regeneration was observed in all basal explants 6 days after meristem excision, followed by growth of the regenerated regions (Figures S6F, S6I, S6L, and S6O). By contrast, in DEX-treated plants, regeneration was absent in some basal explants and considerably reduced in others at the same time point (Figures S6G, S6H, S6M, and S6N). Fourteen days after meristem excision, the areas of regenerated regions in DEX-treated plants had expanded but remained smaller than those in mock-treated plants and lacked normal tissue differentiation (Figures S6I, S6J, S6O, and S6P). Scanning electron microscopy revealed that numerous gemma-like structures with single-layered peripheral regions had formed in the regenerated regions of DEX-treated plants (Figures S6K, S6Q, and S6R). These findings suggest that residual MpGCAM1-specific functions for gemma formation, acting through its downstream targets distinct from those shared with MpGC1L, may underlie this phenotype.

To further examine the roles of downstream genes of MpGC1L in regeneration, we introduced the Mp*EFpro:*Mp*GC1L-SRDX-GR* construct into the Mp*gcam1^ko^* mutant background (Figure 4A). To avoid activating MpGC1L-SRDX during vegetative growth, we treated the plants with DEX 1 day before meristem excision. In mock-treated plants, normal regeneration was observed in basal explants (Figures 4B, 4D, 4F, and 4H), and apical explants grew vigorously (Figures S7B, S7D, S7F, and S7H). By contrast, DEX-treated plants showed severely inhibited regeneration, and no gemma-like structures were observed at the regeneration site, unlike in the WT background (Figures 4C, 4E, 4G, 4I, 4J, and 4K). The growth of apical explants was also inhibited (Figure S7), suggesting that MpGC1L-SRDX activity affects meristem maintenance. These results indicate that the downstream genes of MpGC1L play critical roles in regeneration. Moreover, considering the findings from single and double mutants of Mp*GC1L* and Mp*GCAM1*, it is likely that MpGC1L and MpGCAM1 share a set of downstream genes that are essential for regeneration and may also contribute to meristem maintenance, while MpGCAM1 regulates a specific set of genes for gemma formation.

**Figure 4.**
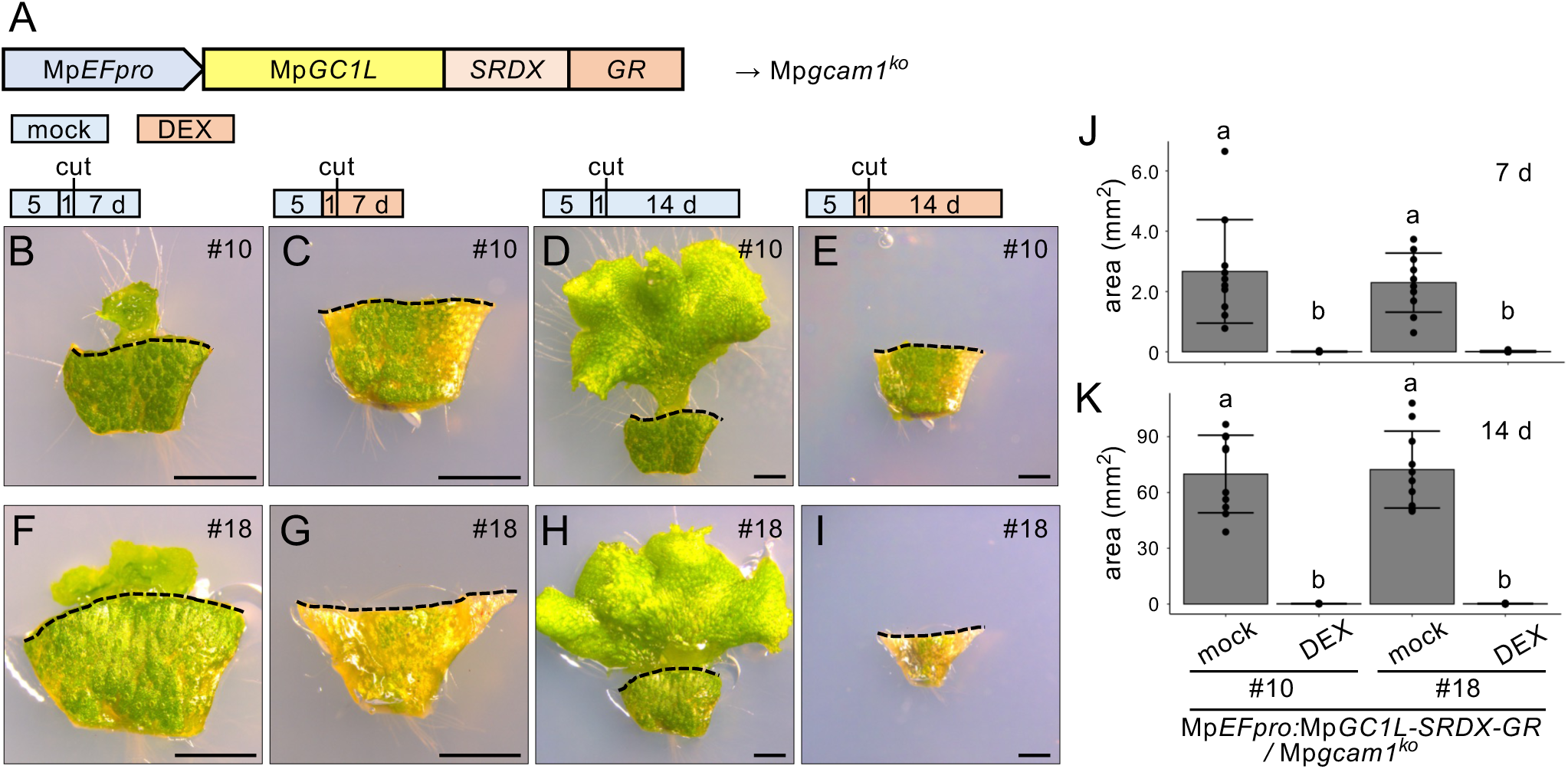
The effects of introducing MpGC1L-SRDX-GR on the regeneration of Mp*gcam1^ko^* plants. (A) Schematic representation of the Mp*EFpro:*Mp*GC1L-SRDX-GR* construct. (B-I) The tips of *pro*Mp*EF:*Mp*GC1L-SRDX-GR* transgenic plants were grown on control medium for 5 days, transferred to control (B, D, F, and H) or DEX (C, E, G, and I) medium, cultured for one day, and their apical meristems removed. The basal explants were treated with mock (B, D, F, and H) or DEX (C, E, G, and I) medium for 7 d (B, C, F, and G) or 14 d (D, E, H, and I). Mp*EFpro:*Mp*GC1L-SRDX-GR /* Mp*gcam1^ko^* #10 (B-E) and #18 (F-I). The bars above the photographs show the DEX treatment conditions. Blue bars, mock treatment; red bars, DEX treatment. The dotted lines show the border between the basal explant and regenerated region. Scale bars = 2 mm. (J and K) Average area of the regenerated region at 7 d (J) and 14 d (K) after excision. Different letters above the bars indicate significant differences at *P* < 0.05 by Tukey–Kramer test. *n* = 10.

### Exploring genes that regulate thallus regeneration downstream of MpGC1L and MpGCAM1

To investigate the genes regulated by MpGC1L and MpGCAM1 during thallus regeneration, we performed RNA-seq analysis using WT and Mp*gc1l^ge^*Mp*gcam1^ko^* double mutant plants. After removing the thallus tips, we cultured the basal explants for 0, 6, or 24 hours. First, we compared WT samples across the time points to identify differentially expressed genes (DEGs) associated with thallus regeneration, identifying a total of 3,955 DEGs. To classify their expression patterns, we conducted k-means clustering analysis. After testing multiple numbers of clusters, we determined that eight clusters provided the clearest separation of expression profiles (Figure 5A).

**Figure 5.**
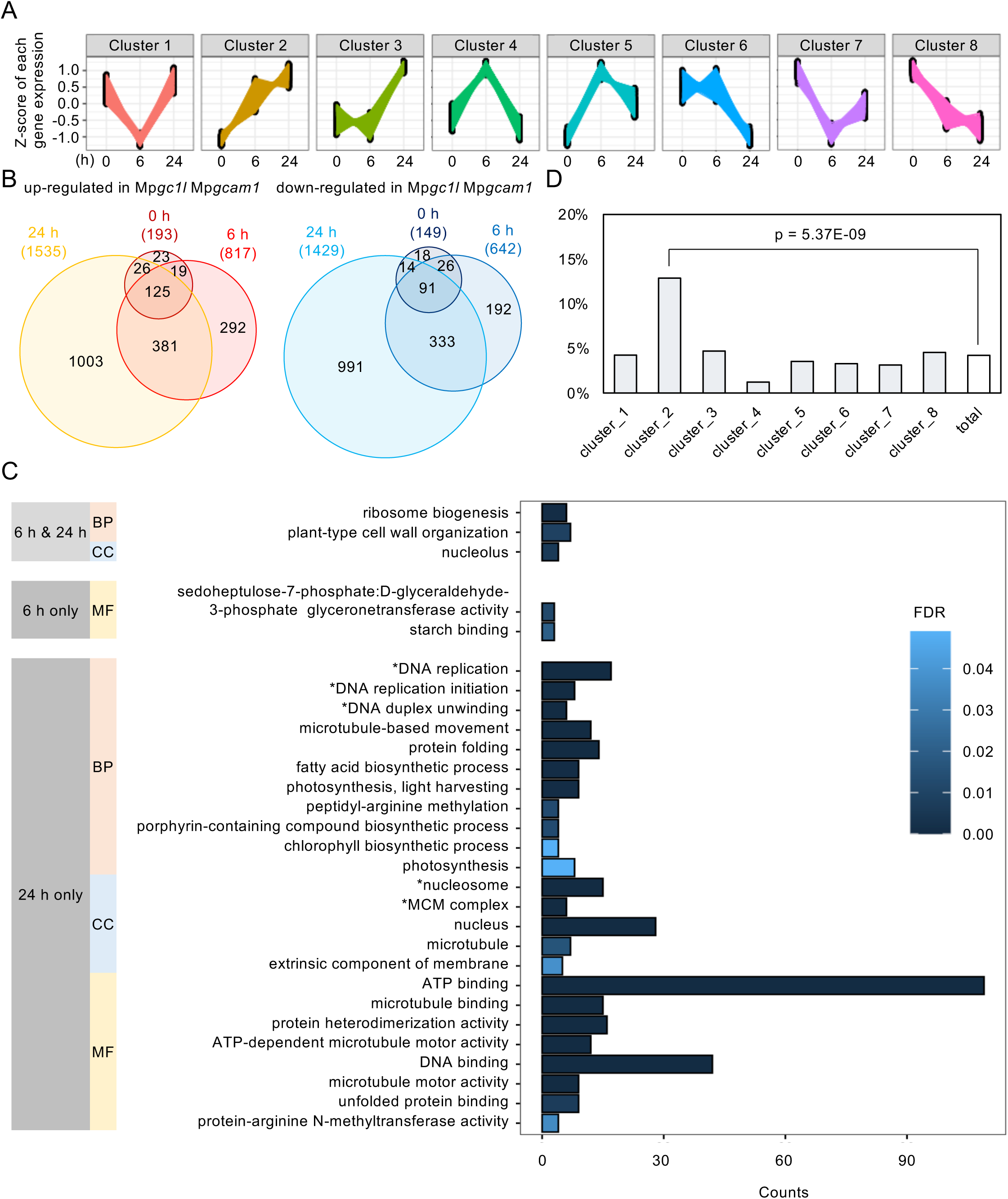
RNA-seq to identify downstream genes of MpGC1L and MpGCAM1 during thallus regeneration. (A) K-means clustering using 3,955 DEGs detected during thallus regeneration in the WT. (B) Venn diagrams of the number of up- and downregulated genes in the Mp*gc1l* Mp*gcam1* double mutant compared to the WT. The number of genes for each fraction is indicated. (C) GO enrichment analysis of downregulated genes in Mp*gc1l* Mp*gcam1.* 333 genes downregulated at 6 and 24 h but not at 0 h, 192 genes downregulated only at 6 h, and 991 genes downregulated only at 24 h were analyzed. Enriched GO terms with FDR < 0.05 are shown. BP: Biological Processes, CC: Cellular Component, MF: Molecular Function. Asterisks show GO terms associated with the cell cycle. (D) Distribution of the 333 downregulated genes in Mp*gc1l* Mp*gcam1* (6 h and 24 h) across the eight clusters defined in the experiment shown in (A). *P*-value between cluster 2 and the total by Fisher’s exact test adjusted by Holm’s method is indicated.

Next, we compared DEGs between WT and Mp*gc1l^ge^* Mp*gcam1^ko^*samples at each time point to identify the genes whose expression depends on MpGC1L and MpGCAM1 during regeneration. We detected 342 DEGs (193 upregulated, 149 downregulated) at 0 hours, 1,459 DEGs (817 upregulated, 642 downregulated) at 6 hours, and 2,964 DEGs (1,535 upregulated, 1,429 downregulated) at 24 hours of culture (Figure 5B). Notably, more than half of upregulated and downregulated genes at 6 hours overlapped with those at later time points (Figure 5B). Chimera-repressor experiments suggest that MpGC1L and MpGCAM1 act as transcriptional activators during thallus regeneration (Figure 4). Therefore, we focused on the genes that were downregulated in the Mp*gc1l^ge^* Mp*gcam1^ko^* double mutant following meristem excision.

Gene ontology (GO) enrichment analysis revealed that GO terms related to ribosome biogenesis were significantly enriched among the 333 genes downregulated in the Mp*gc1l^ge^* Mp*gcam1^ko^* double mutant at both 6 and 24 hours of culture (Figures 5B and 5C). Within this group, genes belonging to cluster_2 in the WT k-means clustering analysis were significantly overrepresented (49 out of 333 genes; Figure 5D). In other words, many genes that were downregulated during early regeneration in Mp*gc1l^ge^* Mp*gcam1 ^ko^* were upregulated during thallus regeneration in the WT (Figure 5A).

Previous studies have shown that ribosome-related genes were enriched among upregulated DEGs during regeneration from wounded tissues or protoplasts in *M. polymorpha*, *A. thaliana* and the moss *Physcomitrium patens* ^19,33–36^. Remarkably, the best BLAST hits of 38 genes (78% of the 49 downregulated DEGs in the Mp*gc1l^ge^* Mp*gcam1^ko^* double mutant at both 6 and 24 hours in cluster_2) in *A. thaliana* were also gradually upregulated within 24 hours during wound-induced callus formation (cluster 3 in the analysis by ^33^; Table S1).

In addition to ribosome biogenesis, GO terms related to the cell cycle were significantly enriched among the 991 genes that were downregulated exclusively at 24 hours in Mp*gc1l^ge^* Mp*gcam1^ko^* (Figure 5C). This finding suggests that MpGC1L and MpGCAM1 activity promotes cell division during regeneration.

### Mp*GC1L* regulates regeneration independently of pathways involving Mp*ERF15* and Mp*LAXR*

Like *GC1L*, Mp*ERF15* is known to be upregulated after incision in *M. polymorpha* ^21^. To investigate the relationship between Mp*GC1L* and Mp*ERF15*, we first examined Mp*ERF15* expression using RNA-seq data from the Mp*gc1l^ge^* Mp*gcam1^ko^*double mutant. Mp*ERF15* was upregulated after apical meristem excision in Mp*gc1l^ge^* Mp*gcam1^ko^*, similar to the WT (Figure 6A). Likewise, OPDA/dn-OPDA biosynthesis genes, including Mp*LOX1*, Mp*LOX4*, Mp*AOS2* and Mp*AOC*, which are known targets of MpERF15 during regeneration ^21^, were also upregulated after apical meristem excision in Mp*gc1l^ge^* Mp*gcam1^ko^* (Figures S8A-S8D). These results indicate that MpGC1L and MpGCAM1 do not regulate the expression of Mp*ERF15* or its downstream OPDA/dn-OPDA biosynthetic genes.

**Figure 6.**
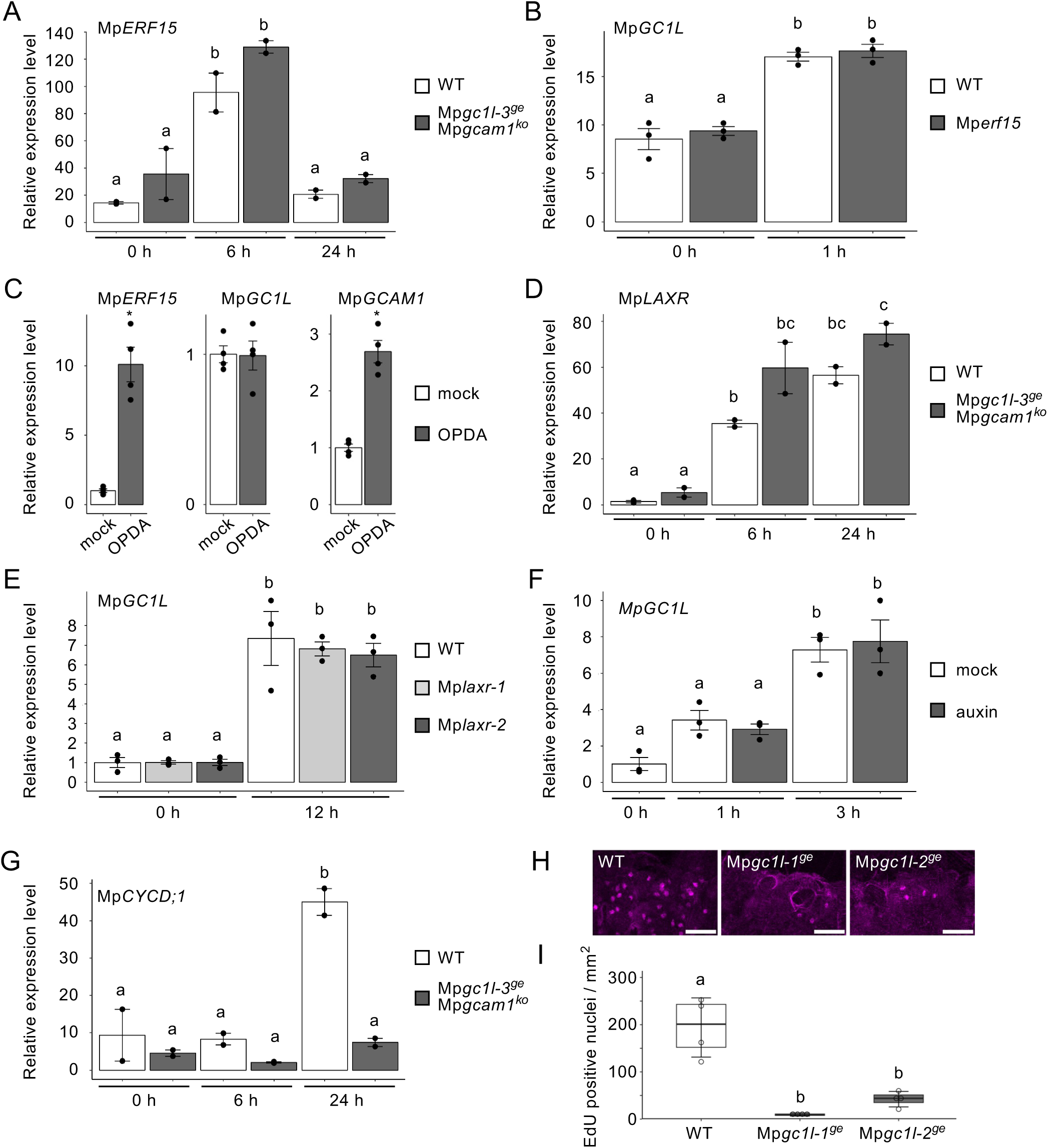
The relationships of MpGC1L and MpGCAM1 with known regeneration pathways. (A-G) Expression analysis of genes involved in regeneration. Relative expression levels of Mp*ERF15* (A), Mp*LAXR* (D), and Mp*CYCD;1* (G) in basal explants after removing the apical meristem based on RNA-seq datasets from WT and Mp*gc1l-3^ge^ gcam1^ko^* plants. Relative expression levels of Mp*GC1L* after wounding based on RNA-seq datasets from WT and Mp*erf15* plants (Liang *et al*., 2022) (B). Relative expression levels of Mp*ERF15*, Mp*GC1L*, and Mp*GCAM1* in gemmalings with or without OPDA treatment, as determined by RT-qPCR (C). Relative expression levels of Mp*GC1L* in basal explants after removing the apical meristem, as determined by RT-qPCR, in WT and Mp*laxr* plants (E) and in WT plants with or without auxin treatment (F). Mp*APT* (C) or Mp*EF1α* (E and F) were used for normalization. Data are mean ± SE. Different letters above the bars indicate significant differences at *p* < 0.05 by Tukey–Kramer test (A, B, D, E, F and G). Asterisk indicates significant difference compared with mock sample at *P* < 0.01 by Welch’s t test (C). Dots indicate biological replicates [*n* = 2 in (A, D and G); *n* = 3 in (B, E and F); *n* = 4 in (C)]. (H and I) EdU incorporation assay to visualize S-phase progression in WT and Mp*gc1l* single mutants. Maximum-projection of confocal images of basal explants labeled with EdU visualized with Alexa Fluor 555 (H). Scale bars = 50 μm. Number of EdU-positive nuclei per area (I). Dots indicate each data point. The box plot shows the interquartile range, spanning from the first quartile to the third quartile. The horizontal line within the box marks the median (second quartile). The whiskers indicate ± SE. Different letters indicate a significant difference (one-way ANOVA; Tukey’s HSD, *p* < 0.05). *n* = 4.

**Figure 7.**
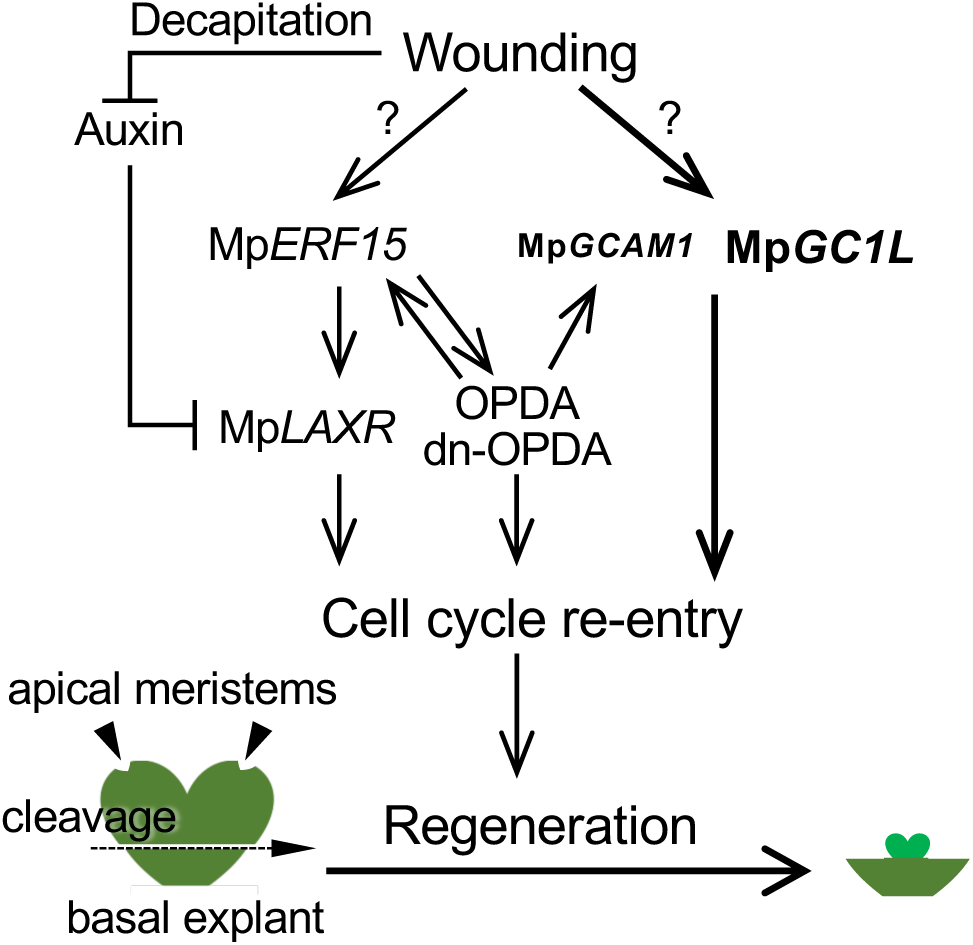
Model of the regulation of regeneration in *M. polymorpha*. In *M. polymorpha*, regeneration occurs from the apical side of the basal explant when the apical meristem is removed even in the absence of plant hormone treatment. MpGC1L plays a major role in regeneration by promoting cell-cycle re-entry, whereas MpGCAM1, which is essential for vegetative reproduction, contributes to regeneration in a more limited capacity. Mp*GC1L* and Mp*GCAM1* function in a separate pathway from Mp*ERF15* (involved in OPDA and dn-OPDA biosynthesis) and Mp*LAXR* (which is induced when auxin levels decrease upon meristem excision [decapitation]).

Next, we analyzed Mp*GC1L* expression in the Mp*erf15* mutant using previously published RNA-seq datasets ^21^. Mp*GC1L* was upregulated 1 hour after wounding in both WT and Mp*erf15* (Figure 6B). Since OPDA biosynthesis and Mp*ERF15* form a positive feedback loop, in which Mp*ERF15* expression is upregulated in response to OPDA treatment ^21^, we examined whether Mp*GC1L* is regulated in a similar manner. Consistent with previous findings, Mp*ERF15* was significantly upregulated by OPDA treatment, while Mp*GC1L* expression was unaffected (Figure 6C). Mp*GCAM1* was also upregulated by OPDA, although to a much lesser extent than Mp*ERF15* (Figure 6C).

We then investigated the relationships of Mp*GC1L* and Mp*GCAM1* with Mp*LAXR*, whose expression is regulated by auxin and MpERF15 ^19,21^. In the RNA-seq datasets of the Mp*gc1l^ge^*Mp*gcam1^ko^* double mutant, Mp*LAXR* was upregulated after excision, similar to the WT (Figure 6D). The expression levels of Mp*GC1L* and Mp*GCAM1* after excision also increased to comparable levels in both Mp*laxr* and WT plants (Figures 6E and S8E). Furthermore, auxin treatment, which suppresses Mp*LAXR* expression ^19^, did not affect the upregulation of Mp*GC1L* or Mp*GCAM1* in WT plants (Figures 6F and S8F).

Finally, we examined the roles of MpGC1L and MpGCAM1 in regulating the cell cycle, as suggested by GO analysis and the previous finding that cell-cycle progression is also regulated by both MpERF15 and MpLAXR ^19,21^. Mp*CYCD;1*, encoding a cell-cycle regulator, was upregulated 24 hours after excision in WT plants but showed little to no activation in Mp*gc1l^ge^* Mp*gcam1^ko^* double mutants (Figure 6G). Additionally, a 5-ethynyl-2’-deoxyuridine (EdU) assay revealed significantly fewer cells undergoing DNA replication in basal explants after meristem excision in Mp*gc1l^ge^* compared to WT plants (Figures 6H, 6I and S9). These results demonstrate that MpGC1L functions independently of pathways involving MpERF15 and MpLAXR during regeneration. Nevertheless, MpGC1L, in cooperation with MpGCAM1, and together with MpERF15 and MpLAXR, acts as a central regulator that drives cell-cycle re-entry, a critical step in regeneration.

## Discussion

We previously identified MpGCAM1 as an essential regulator of vegetative reproduction, gemma and gemma cup formation in *M. polymorpha* ^26^. To date, as in *A. thaliana*, AP2/ERF-type transcription factors, MpLAXR and MpERF15 have also been shown to play important roles in regeneration in *M. polymorpha*. In the present study, we demonstrated that *MpGC1L*, the closest paralog of *MpGCAM1*, a non-AP2/ERF-type transcription factor, is rapidly upregulated in *M. polymorpha* at the regeneration site immediately after removing the apical meristem from thalli. Although the Mp*gc1l^ge^*mutant exhibited a relatively mild yet significant delay in regeneration compared to the WT (Figure 3), EdU assays revealed a clear reduction in the number of cells re-entering the cell cycle (Figures 6H and 6I), indicating that MpGC1L plays a critical role in promoting regeneration. The regeneration phenotype of the Mp*gcam1^ko^* mutant was similar to that of the WT, but the Mp*gc1l^ge^* Mp*gcam1^ko^* double mutants showed a significantly more severe impairment than the Mp*gc1l^ge^* single mutants (Figure 3). These findings suggest that MpGCAM1 acts in support of MpGC1L during regeneration. However, Mp*GC1L* and Mp*GCAM1* displayed differences in the timing of their upregulation following wounding (Figures 1A and S5) and in their responses to OPDA treatment (Figure 6C). These observations suggest that, although MpGC1L and MpGCAM1 function cooperatively, their expression is regulated through distinct regulatory mechanisms.

In DEX-treated Mp*EFpro:*Mp*GC1L-SRDX-GR* plants, where MpGC1L-SRDX is thought to function as a dominant repressor, regeneration was severely inhibited (Figure 4), strongly suggesting that MpGC1L acts as a transcriptional activator during regeneration. The induction of Mp*EFpro:*Mp*GC1L-SRDX-GR* had different effects on WT versus Mp*gcam1^ko^* plants. One possible explanation is that MpGC1L and MpGCAM1 may share downstream targets involved in regeneration, whereas gemma formation depends on MpGCAM1-specific targets (Figures 4 and S6). In DEX-treated Mp*EFpro:*Mp*GC1L-SRDX-GR* plants in the WT background, the expression of regeneration-related downstream genes was likely repressed, while only MpGCAM1-specific targets remained active, resulting in the formation of gemma-like structures. On the other hand, DEX-treated Mp*EFpro:*Mp*GC1L-SRDX-GR* plants in the Mp*gcam1^ko^*background exhibited severe defects not only in regeneration but also in meristem maintenance (Figures 4 and S7). This may be due to the ectopic overexpression of Mp*GC1L-SRDX-GR* in the meristem region driven by the Mp*EF1*α promoter ^37^. However, the observation of occasional meristem abortion in Mp*gc1l^ge^*Mp*gcam1^ko^* double mutants (Figure S4) suggests that MpGC1L and MpGCAM1 might also contribute to meristem maintenance under certain conditions. Further studies are needed to elucidate the specific roles of MpGC1L and MpGCAM1 in meristem maintenance.

Our findings demonstrate that MpGC1L, with support from MpGCAM1, operates through a pathway distinct from that of MpERF15, which is involved in OPDA and dn-OPDA biosynthesis, and Mp*LAXR*, whose expression is regulated by auxin levels. Both Mp*ERF15* and its downstream targets were upregulated in the Mp*gc1l^ge^* Mp*gcam1^ko^*double mutant, as observed in the WT after meristem excision (Figures 6 and S8). This indicates that MpGC1L and MpGCAM1 are not part of the MpERF15-mediated OPDA and dn-OPDA biosynthesis pathway.

The elevated expression of both Mp*ERF15* and Mp*GC1L* immediately after meristem excision suggests that their induction is triggered by wounding signals via mechanisms that remain to be elucidated. Mp*GCAM1* appears to play a minor role compared to Mp*GC1L*, with its expression regulated in part by the MpERF15/OPDA pathway. Perhaps MpGCAM1 mediates coordination between these two independent pathways. One key regulator of regeneration, Mp*LAXR*, is normally repressed by auxin coming from the apical meristem and is derepressed when the auxin supply is interrupted by meristem removal ^19^. The observation that Mp*GC1L* and Mp*GCAM1* expression levels remained elevated even after auxin treatment suggests that they are regulated by a different signaling pathway.

Regeneration was significantly impaired in the Mp*gc1l^ge^*Mp*gcam1^ko^* double mutants, and *GC1L* was rapidly upregulated after excision of the apical meristem, indicating that MpGC1L is a key regulator that initiates the wound-induced regeneration program. The residual regeneration observed in the double mutant suggests that MpGC1L/MpGCAM1 operate partially in parallel with the MpERF15 and MpLAXR pathways during regeneration. Further studies are needed to clarify the interactions among these pathways and to uncover the precise molecular mechanisms underlying their coordination during regeneration.

In *A. thaliana*, wounding induces cytokinin biosynthesis, and cytokinin signaling is required to activate the cell cycle during regeneration by promoting the expression of cell-cycle-related genes ^33,38^. In this study, we demonstrated that MpGC1L and MpGCAM1 are involved in upregulating cell-cycle-related genes during regeneration (Figures 5C and 6G-6I), which is consistent with the formation of undifferentiated cell masses observed upon ectopic overexpression of these genes (Figure 2) ^26^. A recent study also suggested that Mp*GCAM1* is upregulated by cytokinin biosynthesis and signaling in *M. polymorpha* ^39,40^. However, cytokinin accumulation does not increase at the regenerating site of the basal explant in *M. polymorpha* ^19^. Moreover, regeneration occurs even in the Mp*rrb* mutant, which lacks the only type-B response regulator (RR) responsible for cytokinin response in *M. polymorpha* ^41^. Since type-B RRs are required for cytokinin signal transduction, these findings suggest that cytokinin signaling is not essential for regeneration in *M. polymorpha*. Nevertheless, it is possible that cytokinin signaling contributes to regeneration by upregulating Mp*GCAM1*.

GO analysis showed that MpGC1L and MpGCAM1 regulate the expression of ribosome-related genes during regeneration (Figure 5C). Notably, ribosome-related genes are also upregulated in *A. thaliana* and *P. patens* during regeneration ^33–36^. This suggests that MpGC1L and MpGCAM1 may play a conserved role in activating a core set of genes required for plant regeneration across species. Further investigation of the molecular mechanisms underlying regeneration in *M. polymorpha*, particularly the roles of MpGC1L and MpGCAM1, and comparative studies with other plant species will enhance our understanding of the diversity of molecular processes involved in plant regeneration. Such studies may also help explain the high regeneration efficiency observed in bryophytes.

## Methods

### Plant material and growth conditions

The WT plant used in this study was Takaragaike-1 (Tak-1) ^42^. The Mp*gcam1^ko^*, Mp*laxr-1* and Mp*laxr-2* lines were previously established ^19,26^. All plants were cultured on half-strength Gamborg’s B5 medium ^43^ containing 1% agar under 45-55 μmol photons m^-2^ s^-1^ continuous white light with a cold cathode fluorescent lamp (OPT-40C-N-L; Optrom) or light-emitting diode (VGL-1200W; SYNERGYTEC) at 22℃.

### Generation of transgenic lines

To construct Mp*GC1Lpro:ELUC*, the Mp*GC1L* genomic region, including a 4819-bp fragment upstream of the 11^th^ Asp codon, was amplified from Tak-1 genomic DNA by PCR using KOD FX-Neo (TOYOBO) with the primer set proMpGC1L-pro-L1/proMpGC1L-pro-R2 (Table S2) and cloned into the pENTR/D-TOPO entry vector (Thermo Fisher Scientific). The Mp*GC1L* promoter was then transferred into pMpGWB131 ^44^ using Gateway™ LR Clonase™ II Enzyme mix (Thermo Fisher Scientific) to fuse with *ELUC*.

A DNA fragment encoding the Mp*GC1L* coding sequence (CDS), flanked by attL1 and attL2 sequences was synthesized by Genewiz. To generate Mp*EFpro:* Mp*GC1L-GR*, the Mp*GC1L* CDS flanked by attL1 and attL2 was introduced into the Gateway-compatible binary vectors MpGWB113 via LR reaction ^44^. To generate Mp*EFpro:* Mp*GC1L-SRDX-GR*, the fragment was introduced into the Gateway-compatible binary vectors MpGWB121 and MpGWB321 ^44^, via LR reaction for transformation of Tak-1 and Mp*gcam1^ko^*, respectively.

Loss-of-function mutants of Mp*GC1L* were generated using the CRISPR/Cas9 system as described previously ^45^. The two target sequences located within the first exon of Mp*GC1L* (Figure S3) were selected. Corresponding synthetic oligonucleotide pairs (Table S2) were annealed and inserted into the pMpGE_En03 entry vector. The resulting gRNA cassettes were cloned into pMpGE011 binary vector. All binary vectors were introduced into regenerating thalli of Tak-1 or Mp*gcam1^ko^*via *Agrobacterium tumefaciens* GV2260 as described previously ^17^.

### Luciferase activity assay

To measure luciferase activity in Mp*GC1Lpro:ELUC* thalli, 10-day-old gemmalings were sprayed with 1 mM D-Luciferin solution (FUJIFILM Wako). After excising thallus tips, the *ELUC* signal was detected by using the ImageQuant LAS 4000 (GE healthcare UK Ltd.) with an exposure time of 15 min. The resulting images were analyzed using ImageJ/FIJI ^46^.

### Regeneration assay

After gemmae or apical tip explants were grown under the standard conditions described above, the apical meristems were removed, and rectangular basal explants were excised as described by Ishida et al ^19^, and cultured under the standard growth conditions for the indicated number of days. For DEX treatment, basal explants or intact plants prior to excision were transferred to solid medium containing 10 μM DEX (Sigma-Aldrich) and cultured.

Photographs of regenerating buds were taken with the stereomicroscope (M205C, Leica Microsystems) or M205 FA (Leica Microsystems). The areas of regeneration region were measured using ImageJ/FIJI ^46^ with the area measurement tool. For scanning electron microscopy (SEM), plant samples were frozen in liquid nitrogen and directly observed using a VHX-D500 microscope (KEYENCE).

### Phenotypic analysis of Mp*EFpro:*Mp*GC1L-GR* plants

Gemmae of Mp*EFpro:*Mp*GC1L-GR* plants were grown on solid medium containing 10 μM DEX for 14 days. The resulting phenotypes were observed using a stereomicroscope, M205 FA (Leica Microsystems). For SEM analysis, plant samples were frozen using liquid nitrogen and directly observed using a VHX-D500 microscope (KEYENCE).

### Quantitative real-time RT-PCR

For OPDA treatment, 10-day-old WT gemmalings were overlaid with a 10 µM OPDA solution (Cayman Chemical Co.) or a mock solution (1% ethanol) so that they were submerged, and incubated for 2 h before whole plants were collected. For auxin treatment, basal explants were treated with 1 µM 1-naphthaleneacetic acid (NAA) and harvested as previously described ^19^. For the other experiments, basal explants of WT, Mp*gc1l-1*, Mp*laxr-1* and Mp*laxr-2* were harvested. Total RNA was extracted from these samples using either the RNeasy plant mini kit (Qiagen) or TRIzol reagent (Thermo Fisher Scientific), followed by DNase treatment with the RNase-Free DNase set (Qiagen). Reverse transcription was performed with an oligo(dT) primer, or a combination of oligo(dT) primer and random hexamer primers using ReverTraAce® or ReverTra Ace® with gDNA Remover (TOYOBO). Quantitative PCR was conducted on a CFX96 Real-Time PCR Detection System (Bio-Rad) using SYBR® Green I Nucleic Gel Stain (Lonza) or with a LightCycler® 96 instrument (Roche) using KOD SYBR® qPCR Mix (TOYOBO). Each sample was measured in triplicate. Expression of Mp*EF1α* or Mp*APT* was used for normalization. Primer sequences are listed in Table S2.

### Transcriptome analysis

Total RNA was extracted from WT and Mp*gc1l-3^ge^* Mp*gcam1^ko^*basal explants using the RNeasy plant mini kit (Qiagen) with on-column DNase treatment (RNase-Free DNase set, Qiagen). Library preparation and RNA-seq analysis were outsourced to Genewiz (https://www.genewiz.com/). Libraries were prepared using NEBNext Ultra II RNA Library Prep Kit for Illumina (New England Biolab) and sequenced as paired-end reads on an Hiseq4000 (Illumina).

Raw read quality was assessed using FastQC (www.bioinformatics.babraham.ac.uk/projects/fastqc). Illumina adapters at the ends of the paired reads were cleaned up using Cutadapt 2.1 ^47^. The cleaned FASTQ reads were mapped onto the Marchantia polymorpha genome (v6.1 accessed through Marpolbase: https://marchantia.info/) using HISAT2 v2.1.0 ^48^ with default parameters. Post-processing of SAM/BAM files was performed using SAMTOOLS v1.9. ^49^. GFF file was converted into GTF format using gffread embedded in cufflinks ^50^, and then FeatureCounts v1.6.4 ^51^ was used to count the raw reads corresponding to each gene. EdgeR ^52^ was used to normalize the raw counts and perform the differential expression analysis (Padj < 0.01). K-means clustering was performed by converting normalized read count to Z-score using RStudio (http://www.rstudio.com/). GO enrichment analysis was performed using PlantRegMap by converting gene ID from v6.1 to v3.1 ^53^. All the sequenced raw reads were deposited in DDBJ under the BioProjrct number PRJDB39661. Previously reported RNA-seq data sets using Mp*erg15* and WT as a control were downloaded (SRR18054067-SRR18054078 ^21^) from SRA (https://www.ncbi.nlm.nih.gov/sra). RNA-seq read files were cleaned using Fastp v0.20.0 ^54^ and the read were mapped to the *M. polymorpha* assemble genome (v.6.1) using HISAT2 v2.2.1 ^48^. The mapped reads were sorted and indexed using SAMtools v1.10 ^49^ and counted using StringTie v.2.1.3 ^55^ with Pyson package ‘prepDE.py’ using MpTak_v6.1r1.gff (MarpolBase: https://marchantia.info) as a reference.

### Visualization of S-phase cells

To assess the cell cycle re-entry after the apical meristem excision, S-phase nuclei were visualized using EdU staining as described previously with slight modifications ^56^. The apical meristems were removed from 10-day-old plants after gemma germination. These plants were then cultured for 44 hours and subsequently submerged in half strength Gamborg’s B5 liquid medium with 20 μM EdU for 4 h under the standard growth conditions described above. Samples were fixed in FAA solution [2.5% Formalin, 2.5 % acetic acid, and 50 % ethanol]. EdU incorporation was detected with Alexa Fluor 555-azide following the manufacturer’s protocol (Click-iT™ Plus EdU Cell Proliferation Kit for Imaging, Alexa Fluor™ 555 dye; Thermo Fisher Scientific). For clearing and cell wall staining, samples were treated for 4 hours with the ClearSee solution [10% (w/v) xylitol, 15% (w/v) sodium deoxycholate, and 25% (w/v) urea] with 0.02% (v/v) SCRI Renaissance 2200 (Renaissance Chemicals, Selby, UK) ^57^. After replacement with fresh ClearSee solution, samples were incubated for at least three additional days before microscopic observation. Samples were mounted in ClearSee solution between two cover glasses and observed using an inverted microscope IX70 (Evident) equipped with a confocal laser scanning system FV1000 (Evident). Thirty-one confocal images were acquired at 5-µm intervals, with excitation (559 nm) and emission (603 nm) settings for Alexa Fluor 555 dye, and excitation (405 nm) and emission (461 nm) settings for SCRI Renaissance 2200 dye. Z-stacked images were reconstructed and analyzed using ImageJ/FIJI program ^46^.

## Acknowledgments

The authors thank Sakiko Ishida for identifying the marked induction of MpGC1L during regeneration from thallus fragments and for providing the initial data that motivated this study and performing gene expression analysis, and Chiho Hirata and Eri Okada for technical assistance. The research was funded by Japan Society for the Promotion of Science (JSPS) KAKENHI, Grant-in-Aid for Early-Career Scientists (19K16167 to Y.Y.), Grant-in-Aid for Transformative Research Areas (23H04744 and 25H01820 to Y.Y.), Grant-in-Aid for Scientific Research (B) (15H04391 and 19H03247 to K.I.), Grant-in-Aid for Scientific Research on Priority Areas (25119711, 15H01233, and 17H06472 to K.I.), JSPS the Program for Forming Japan’s Peak Research Universities (J-PEAKS; grant no JPJS00420230009 to K.I.), Grant-in-Aid for Research Activity Start-up (19K23751 to H.K.), Grant-in-Aid for Early-Career Scientists (21K15125 to H.K.), Grant-in-Aid for JSPS Fellows (21J40092 for Y.S.), Grant-in-Aid for Scientific Research (C) (25K09689 for Y.S.). Grant-in-Aid for Scientific Research (B) (JP23H02505 to R.N.) and Grant-in-Aid for Scientific Research on Priority Areas (JP18H04836, JP20H04884 and JP22H04733 to R.N.). We also thank the Research Facility Center for Science and Technology of Kobe University for supporting our experiments. It was also supported by the JST FOREST Program (JPMJFR2256 to Y.Y.), JST A-STEP Program (JPMJTR25U4 to KI) and JST GteX Program (JPMJGX23B0 to K.I.)

During the preparation of this work, the authors used ChatGPT (OpenAI) for language editing and improvement of English expression. The authors reviewed and edited the output and take full responsibility for the content of the published article.

## References

1. Bowman, J.L., and Eshed, Y. (2000). Formation and maintenance of the shoot apical meristem. Trend in Plant Science 5, 110–115. doi: 10.1016/s1360-1385(00)01569-7.

2. Arnoux-Courseaux, M., and Coudert, Y. (2024). Re-examining meristems through the lens of evo-devo. Trends Plant Sci 29, 413–427. 10.1016/j.tplants.2023.11.003.

3. Cao, X., and Jiao, Y. (2020). Control of cell fate during axillary meristem initiation. Cell Mol Life Sci 77, 2343–2354. 10.1007/s00018-019-03407-8.

4. Ikeuchi, M., Ogawa, Y., Iwase, A., and Sugimoto, K. (2016). Plant regeneration: cellular origins and molecular mechanisms. Development 143, 1442–1451. 10.1242/dev.134668.

5. Williams, L.E. (2021). Genetics of shoot meristem and shoot regeneration. Annu Rev Genet 55, 661–681. 10.1146/annurev-genet-071719-020439.

6. Atta, R., Laurens, L., Boucheron-Dubuisson, E., Guivarc’h, A., Carnero, E., Giraudat-Pautot, V., Rech, P., and Chriqui, D. (2009). Pluripotency of Arabidopsis xylem pericycle underlies shoot regeneration from root and hypocotyl explants grown in vitro. Plant J 57, 626–644. 10.1111/j.1365-313X.2008.03715.x.

7. Kareem, A., Durgaprasad, K., Sugimoto, K., Du, Y., Pulianmackal, A.J., Trivedi, Z.B., Abhayadev, P.V., Pinon, V., Meyerowitz, E.M., Scheres, B., and Prasad, K. (2015). *PLETHORA* genes control regeneration by a two-step mechanism. Curr Biol 25, 1017–1030. 10.1016/j.cub.2015.02.022.

8. Banno, H., Ikeda, Y., Niu, Q.W., and Chua, N.H. (2001). Overexpression of Arabidopsis ESR1 induces initiation of shoot regeneration. Plant Cell 13, 2609–2618. 10.1105/tpc.010234.

9. Gordon, S.P., Heisler, M.G., Reddy, G.V., Ohno, C., Das, P., and Meyerowitz, E.M. (2007). Pattern formation during de novo assembly of the Arabidopsis shoot meristem. Development 134, 3539–3548. 10.1242/dev.010298.

10. Matsuo, N., Makino, M., and Banno, H. (2011). *Arabidopsis ENHANCER OF SHOOT REGENERATION* (*ESR*)1 and *ESR2* regulate in vitro shoot regeneration and their expressions are differentially regulated. Plant Sci. 181, 39–46. 10.1016/j.plantsci.2011.03.007.

11. Ikeuchi, M., Favero, D.S., Sakamoto, Y., Iwase, A., Coleman, D., Rymen, B., and Sugimoto, K. (2019). Molecular mechanisms of plant regeneration. Annu Rev Plant Biol 70, 377–406. 10.1146/annurev-arplant-050718-100434.

12. Zhang, G., Zhao, F., Chen, L., Pan, Y., Sun, L., Bao, N., Zhang, T., Cui, C.X., Qiu, Z., Zhang, Y., et al. (2019). Jasmonate-mediated wound signalling promotes plant regeneration. Nat Plants 5, 491–497. 10.1038/s41477-019-0408-x.

13. Zhou, W., Lozano-Torres, J.L., Blilou, I., Zhang, X., Zhai, Q., Smant, G., Li, C., and Scheres, B. (2019). A jasmonate signaling network activates root stem cells and promotes regeneration. Cell 177, 942–956 e914. 10.1016/j.cell.2019.03.006.

14. Zhang, G., Liu, W., Gu, Z., Wu, S. E Y., Zhou, W., Lin, J., and Xu, L. (2023). Roles of the wound hormone jasmonate in plant regeneration. J Exp Bot 74, 1198–1206. 10.1093/jxb/erab508.

15. Hohe, A., and Reski, R. (2005). From axenic spore germination to molecular farming. One century of bryophyte in vitro culture. Plant Cell Rep 23, 513–521. 10.1007/s00299-004-0894-8.

16. Shimamura, M. (2016). Marchantia polymorpha: taxonomy, phylogeny and morphology of a model system. Plant Cell Physiol 57, 230–256. 10.1093/pcp/pcv192.

17. Kubota, A., Ishizaki, K., Hosaka, M., and Kohchi, T. (2013). Efficient Agrobacterium-mediated transformation of the liverwort Marchantia polymorpha using regenerating thalli. Biosci Biotechnol Biochem 77, 167–172. 10.1271/bbb.120700.

18. Nishihama, R., Ishizaki, K., Hosaka, M., Matsuda, Y., Kubota, A., and Kohchi, T. (2015). Phytochrome-mediated regulation of cell division and growth during regeneration and sporeling development in the liverwort Marchantia polymorpha. J Plant Res 128, 407–421. 10.1007/s10265-015-0724-9.

19. Ishida, S., Suzuki, H., Iwaki, A., Kawamura, S., Yamaoka, S., Kojima, M., Takebayashi, Y., Yamaguchi, K., Shigenobu, S., Sakakibara, H., et al. (2022). Diminished auxin signaling triggers cellular reprogramming by inducing a regeneration factor in the liverwort *Marchantia polymorpha*. Plant Cell Physiol 63, 384–400. 10.1093/pcp/pcac004.

20. Flores-Sandoval, E., Nishihama, R., and Bowman, J.L. (2024). Hormonal and genetic control of pluripotency in bryophyte model systems. Curr Opin Plant Biol 77, 102486. 10.1016/j.pbi.2023.102486.

21. Liang, Y., Heyman, J., Xiang, Y., Vandendriessche, W., Balkan, C., Goeminne, G., and De Veylder, L. (2022). The wound-activated ERF15 transcription factor drives Marchantia polymorpha regeneration by activating an oxylipin biosynthesis feedback loop. Science Advances 8, eabo7737. 10.1126/sciadv.abo7737.

22. Monte, I., Ishida, S., Zamarreno, A.M., Hamberg, M., Franco-Zorrilla, J.M., Garcia-Casado, G., Gouhier-Darimont, C., Reymond, P., Takahashi, K., Garcia-Mina, J.M., et al. (2018). Ligand-receptor co-evolution shaped the jasmonate pathway in land plants. Nat Chem Biol 14, 480–488. 10.1038/s41589-018-0033-4.

23. Kaji, T., Nishizato, Y., Yoshimatsu, H., Yoda, A., Liang, W., Chini, A., Fernandez-Barbero, G., Nozawa, K., Kyozuka, J., Solano, R., and Ueda, M. (2024). Delta(4)-dn-iso-OPDA, a bioactive plant hormone of Marchantia polymorpha. iScience 27, 110191. 10.1016/j.isci.2024.110191.

24. Kaji, T., Nishizato, Y., Matsumoto, K., Okumura, T., and Ueda, M. (2026). Advances in the chemical biology of jasmonates. J Exp Bot. 10.1093/jxb/erag156.

25. Suzuki, H., Harrison, C.J., Shimamura, M., Kohchi, T., and Nishihama, R. (2020). Positional cues regulate dorsal organ formation in the liverwort Marchantia polymorpha. J Plant Res 133, 311–321. 10.1007/s10265-020-01180-5.

26. Yasui, Y., Tsukamoto, S., Sugaya, T., Nishihama, R., Wang, Q., Kato, H., Yamato, K.T., Fukaki, H., Mimura, T., Kubo, H., et al. (2019). GEMMA CUP-ASSOCIATED MYB1, an ortholog of axillary meristem regulators, is essential in vegetative reproduction in *Marchantia polymorpha*. Curr Biol 29, 3987–3995 e3985. 10.1016/j.cub.2019.10.004.

27. Kato, H., Yasui, Y., and Ishizaki, K. (2020). Gemma cup and gemma development in Marchantia polymorpha. New Phytol 228, 459–465. 10.1111/nph.16655.

28. Stracke, R., Werber, M., and Weisshaar, B. (2001). The *R2R3-MYB* gene family in *Arabidopsis thaliana*. Curr Opin Plant Biol 4, 447–456. 10.1016/s1369-5266(00)00199-0.

29. Muller, D., Schmitz, G., and Theres, K. (2006). *Blind* homologous *R2R3 Myb* genes control the pattern of lateral meristem initiation in *Arabidopsis*. Plant Cell 18, 586–597. 10.1105/tpc.105.038745.

30. Kawamura, S., Romani, F., Yagura, M., Mochizuki, T., Sakamoto, M., Yamaoka, S., Nishihama, R., Nakamura, Y., Yamato, K.T., Bowman, J.L., et al. (2022). MarpolBase Expression: A web-based, comprehensive platform for visualization and analysis of transcriptomes in the liverwort *Marchantia polymorpha*. Plant Cell Physiol 63, 1745–1755. 10.1093/pcp/pcac129.

31. Bowman, J.L., Kohchi, T., Yamato, K.T., Jenkins, J., Shu, S., Ishizaki, K., Yamaoka, S., Nishihama, R., Nakamura, Y., Berger, F., et al. (2017). Insights into Land Plant Evolution Garnered from the Marchantia polymorpha Genome. Cell 171, 287–304 e215. 10.1016/j.cell.2017.09.030.

32. Hiratsu, K., Matsui, K., Koyama, T., and Ohme-Takagi, M. (2003). Dominant repression of target genes by chimeric repressors that include the EAR motif, a repression domain, in Arabidopsis. Plant J 34, 733–739. 10.1046/j.1365-313x.2003.01759.x.

33. Ikeuchi, M., Iwase, A., Rymen, B., Lambolez, A., Kojima, M., Takebayashi, Y., Heyman, J., Watanabe, S., Seo, M., De Veylder, L., et al. (2017). Wounding triggers callus formation via dynamic hormonal and transcriptional changes. Plant Physiol 175, 1158–1174. 10.1104/pp.17.01035.

34. Chupeau, M.C., Granier, F., Pichon, O., Renou, J.P., Gaudin, V., and Chupeau, Y. (2013). Characterization of the early events leading to totipotency in an *Arabidopsis* protoplast liquid culture by temporal transcript profiling. Plant Cell 25, 2444–2463. 10.1105/tpc.113.109538.

35. Xiao, L., Zhang, L., Yang, G., Zhu, H., and He, Y. (2012). Transcriptome of protoplasts reprogrammed into stem cells in *Physcomitrella patens*. PLoS One 7, e35961. 10.1371/journal.pone.0035961.

36. Kubo, M., Nishiyama, T., Tamada, Y., Sano, R., Ishikawa, M., Murata, T., Imai, A., Lang, D., Demura, T., Reski, R., and Hasebe, M. (2019). Single-cell transcriptome analysis of Physcomitrella leaf cells during reprogramming using microcapillary manipulation. Nucleic Acids Res 47, 4539–4553. 10.1093/nar/gkz181.

37. Althoff, F., Kopischke, S., Zobell, O., Ide, K., Ishizaki, K., Kohchi, T., and Zachgo, S. (2014). Comparison of the MpEF1alpha and CaMV35 promoters for application in Marchantia polymorpha overexpression studies. Transgenic Res 23, 235–244. 10.1007/s11248-013-9746-z.

38. Iwase, A., Mitsuda, N., Koyama, T., Hiratsu, K., Kojima, M., Arai, T., Inoue, Y., Seki, M., Sakakibara, H., Sugimoto, K., and Ohme-Takagi, M. (2011). The AP2/ERF transcription factor WIND1 controls cell dedifferentiation in Arabidopsis. Curr Biol 21, 508–514. 10.1016/j.cub.2011.02.020.

39. Aki, S.S., Morimoto, T., Ohnishi, T., Oda, A., Kato, H., Ishizaki, K., Nishihama, R., Kohchi, T., and Umeda, M. (2022). R2R3-MYB transcription factor GEMMA CUP-ASSOCIATED MYB1 mediates the cytokinin signal to achieve proper organ development in Marchantia polymorpha. Sci Rep 12, 21123. 10.1038/s41598-022-25684-3.

40. Komatsu, A., Fujibayashi, M., Kumagai, K., Suzuki, H., Hata, Y., Takebayashi, Y., Kojima, M., Sakakibara, H., and Kyozuka, J. (2025). KAI2-dependent signaling controls vegetative reproduction in Marchantia polymorpha through activation of LOG-mediated cytokinin synthesis (14). Nat Commun 16, 1263. 10.1038/s41467-024-55728-3.

41. Bao, H., Sun, R., Iwano, M., Yoshitake, Y., Aki, S.S., Umeda, M., Nishihama, R., Yamaoka, S., and Kohchi, T. (2024). Conserved CKI1-mediated signaling is required for female germline specification in Marchantia polymorpha. Curr Biol 34, 1324–1332 e1326. 10.1016/j.cub.2024.01.013.

42. Ishizaki, K., Chiyoda, S., Yamato, K.T., and Kohchi, T. (2008). Agrobacterium-mediated transformation of the haploid liverwort Marchantia polymorpha L., an emerging model for plant biology. Plant Cell Physiol 49, 1084–1091. 10.1093/pcp/pcn085.

43. Gamborg, O.L., Miller, R.A., and Ojima, K. (1968). Nutrient requirements of suspension cultures of soybean root cells. Experimental Cell Research 50, 151–158. 10.1016/0014-4827(68)90403-5.

44. Ishizaki, K., Nishihama, R., Ueda, M., Inoue, K., Ishida, S., Nishimura, Y., Shikanai, T., and Kohchi, T. (2015). Development of Gateway Binary Vector Series with Four Different Selection Markers for the Liverwort Marchantia polymorpha. PLoS One 10, e0138876. 10.1371/journal.pone.0138876.

45. Sugano, S.S., Nishihama, R., Shirakawa, M., Takagi, J., Matsuda, Y., Ishida, S., Shimada, T., Hara-Nishimura, I., Osakabe, K., and Kohchi, T. (2018). Efficient CRISPR/Cas9-based genome editing and its application to conditional genetic analysis in Marchantia polymorpha. PLoS One 13, e0205117. 10.1371/journal.pone.0205117.

46. Schindelin, J., Arganda-Carreras, I., Frise, E., Kaynig, V., Longair, M., Pietzsch, T., Preibisch, S., Rueden, C., Saalfeld, S., Schmid, B., et al. (2012). Fiji: an open-source platform for biological-image analysis. Nat Methods 9, 676–682. 10.1038/nmeth.2019.

47. Martin, M. (2011). Cutadapt removes adapter sequences from high-throughput sequencing reads. EMBnet. journal 17. 10.14806/ej.17.1.200.

48. Kim, D., Paggi, J.M., Park, C., Bennett, C., and Salzberg, S.L. (2019). Graph-based genome alignment and genotyping with HISAT2 and HISAT-genotype. Nat Biotechnol 37, 907–915. 10.1038/s41587-019-0201-4.

49. Li, H., Handsaker, B., Wysoker, A., Fennell, T., Ruan, J., Homer, N., Marth, G., Abecasis, G., Durbin, R., and Genome Project Data Processing, S. (2009). The Sequence Alignment/Map format and SAMtools. Bioinformatics 25, 2078–2079. 10.1093/bioinformatics/btp352.

50. Trapnell, C., Williams, B.A., Pertea, G., Mortazavi, A., Kwan, G., van Baren, M.J., Salzberg, S.L., Wold, B.J., and Pachter, L. (2010). Transcript assembly and quantification by RNA-Seq reveals unannotated transcripts and isoform switching during cell differentiation. Nat Biotechnol 28, 511–515. 10.1038/nbt.1621.

51. Liao, Y., Smyth, G.K., and Shi, W. (2014). featureCounts: an efficient general purpose program for assigning sequence reads to genomic features. Bioinformatics 30, 923–930. 10.1093/bioinformatics/btt656.

52. Robinson, M.D., McCarthy, D.J., and Smyth, G.K. (2010). edgeR: a Bioconductor package for differential expression analysis of digital gene expression data. Bioinformatics 26, 139–140. 10.1093/bioinformatics/btp616.

53. Tian, F., Yang, D.C., Meng, Y.Q., Jin, J., and Gao, G. (2020). PlantRegMap: charting functional regulatory maps in plants. Nucleic Acids Res 48, D1104–D1113. 10.1093/nar/gkz1020.

54. Chen, S., Zhou, Y., Chen, Y., and Gu, J. (2018). fastp: an ultra-fast all-in-one FASTQ preprocessor. Bioinformatics 34, i884–i890. 10.1093/bioinformatics/bty560.

55. Pertea, M., Kim, D., Pertea, G.M., Leek, J.T., and Salzberg, S.L. (2016). Transcript-level expression analysis of RNA-seq experiments with HISAT, StringTie and Ballgown. Nat Protoc 11, 1650–1667. 10.1038/nprot.2016.095.

56. Furuya, T., Nishihama, R., Ishizaki, K., Kohchi, T., Fukuda, H., and Kondo, Y. (2022). A glycogen synthase kinase 3-like kinase MpGSK regulates cell differentiation in Marchantia polymorpha. Plant Biotechnol (Tokyo) 39, 65–72. 10.5511/plantbiotechnology.21.1219a.

57. Kurihara, D., Mizuta, Y., Sato, Y., and Higashiyama, T. (2015). ClearSee: a rapid optical clearing reagent for whole-plant fluorescence imaging. Development 142, 4168–4179. 10.1242/dev.127613.

