## Supplemental Data (Figures and Tables) for "A paralog of a clonal propagation regulator promotes cell-cycle re-entry during thallus regeneration in *Marchantia polymorpha*"

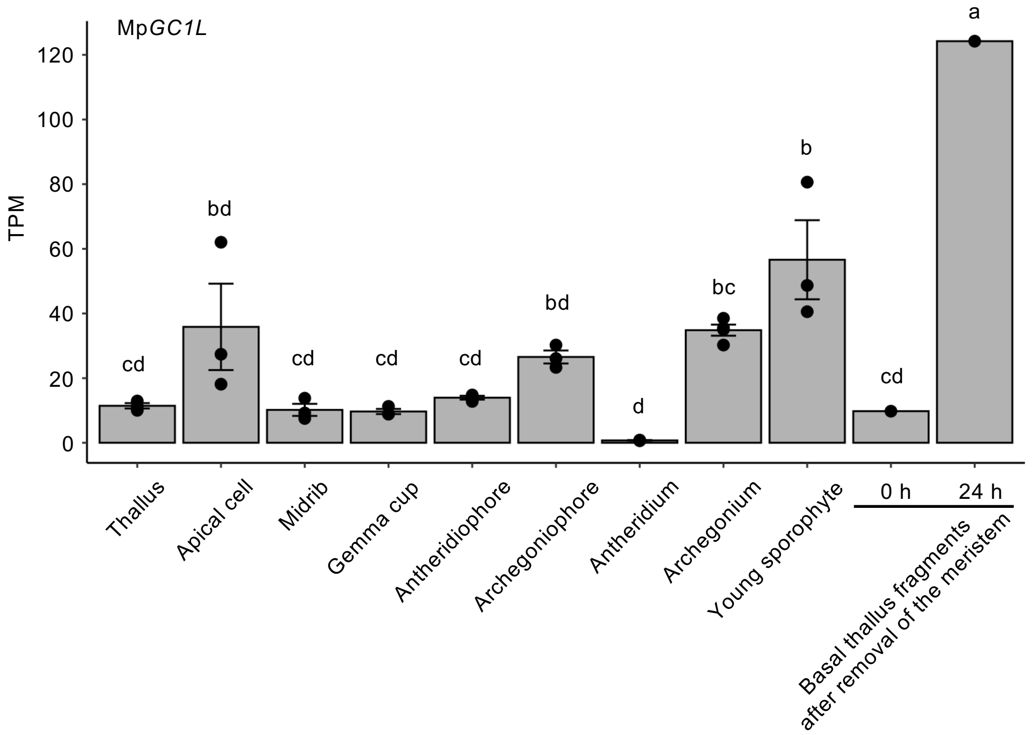


**Figure S1. Mp*GC1L* expression levels in *M. polymorpha* in the MarpolBase Expression database (MBEX) .**

Data are means ± SE. Different letters above the bars indicate significant differences at $p < 0.05$ by Tukey–Kramer test. Dots indicate biological replicates.

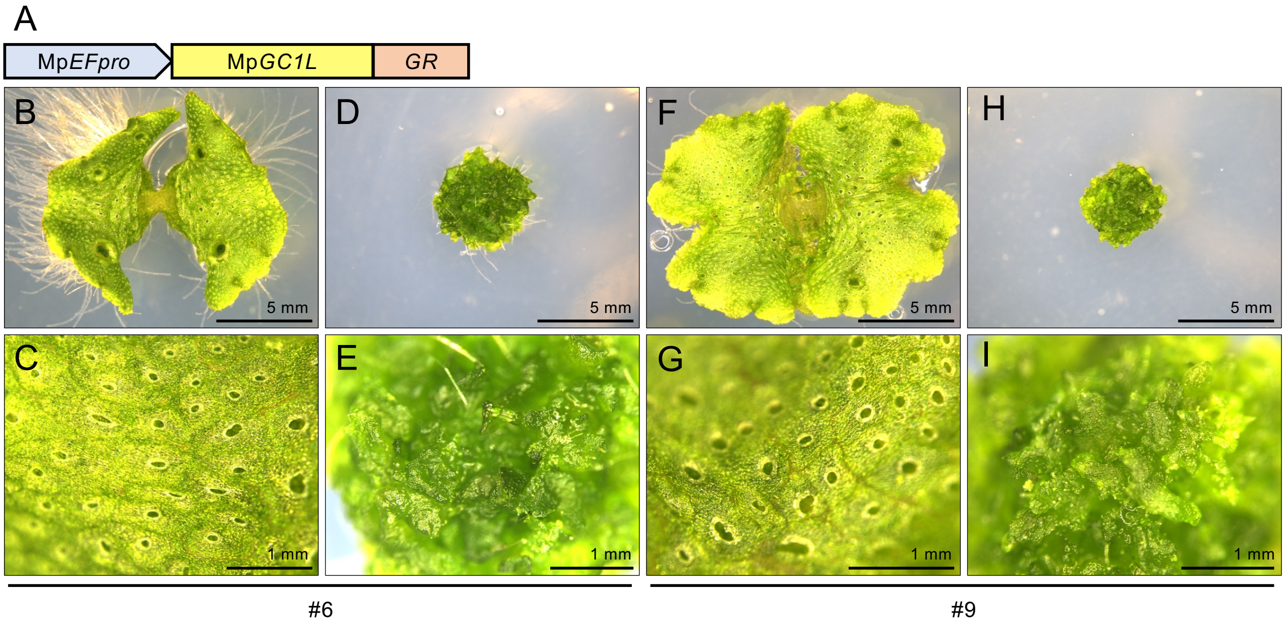


**Figure S2. Inducible ectopic overexpression of MpGC1L function.**
(A) Schematic representation of the Mp*EFpro:*Mp*GC1L-GR* construct. (B-E) Two-week-old Mp*EFpro:*Mp*GC1L-GR* #6 transgenic plants subjected to mock (B and C) or 10 μM DEX treatment (D and E). Images of whole plants (B and D). High-magnification views of the plants (C and E). (F-I) Two-week-old Mp*EFpro:*Mp*GC1L-GR* #9 transgenic plants subjected to mock (F and G) or 10 μM DEX treatment (H and I). Images of whole plants (F and H). High-magnification views of the plants (G and I).

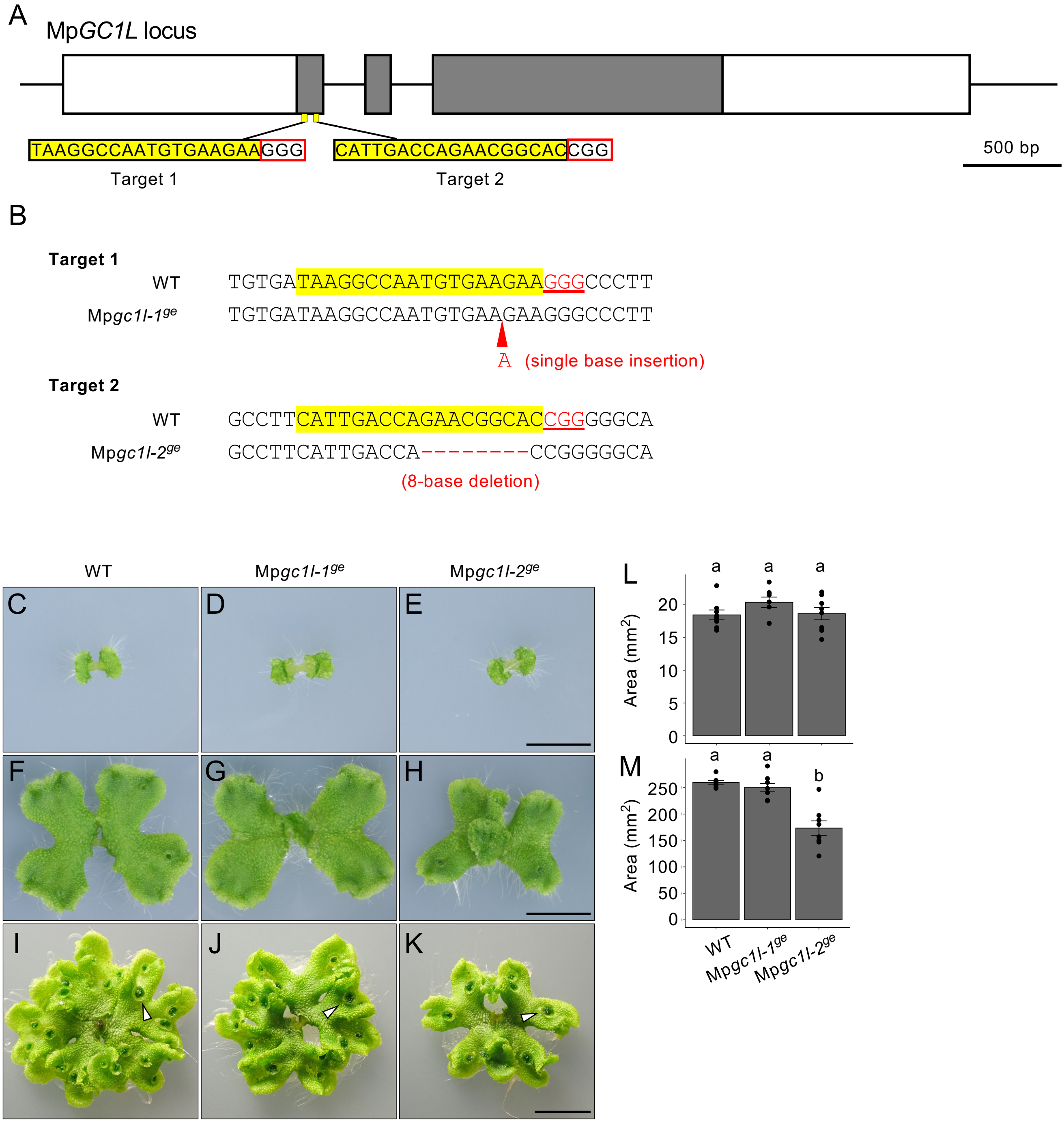


**Figure S3. Sequences of the target sites and phenotypes of Mp*gc1l* single mutants.**
(A) Schematic representation of the two independent target sites and sequences in Mp*GC1L*. (B) Genome sequences of the target sites in the Mp*gc1l*$^{ge}$ mutants in the WT background. (C-K) Thalli of the WT (C, F and I), Mp*gc1l-1*$^{ge}$ mutant (D, G and J) and Mp*gc1l-2*$^{ge}$ mutant (E, H and K) grown from gemma for 7 d (C-E), 14 d (F-H) or 24 d (I-K). Arrowheads indicate representative gemma cup. Bars = 1 cm (L and M) Quantification of thallus area at 7 d or 14 d. Letters above the bars indicate significant differences (Games-Howell test following Welch's one-way ANOVA, $p < 0.05$). n = 8.

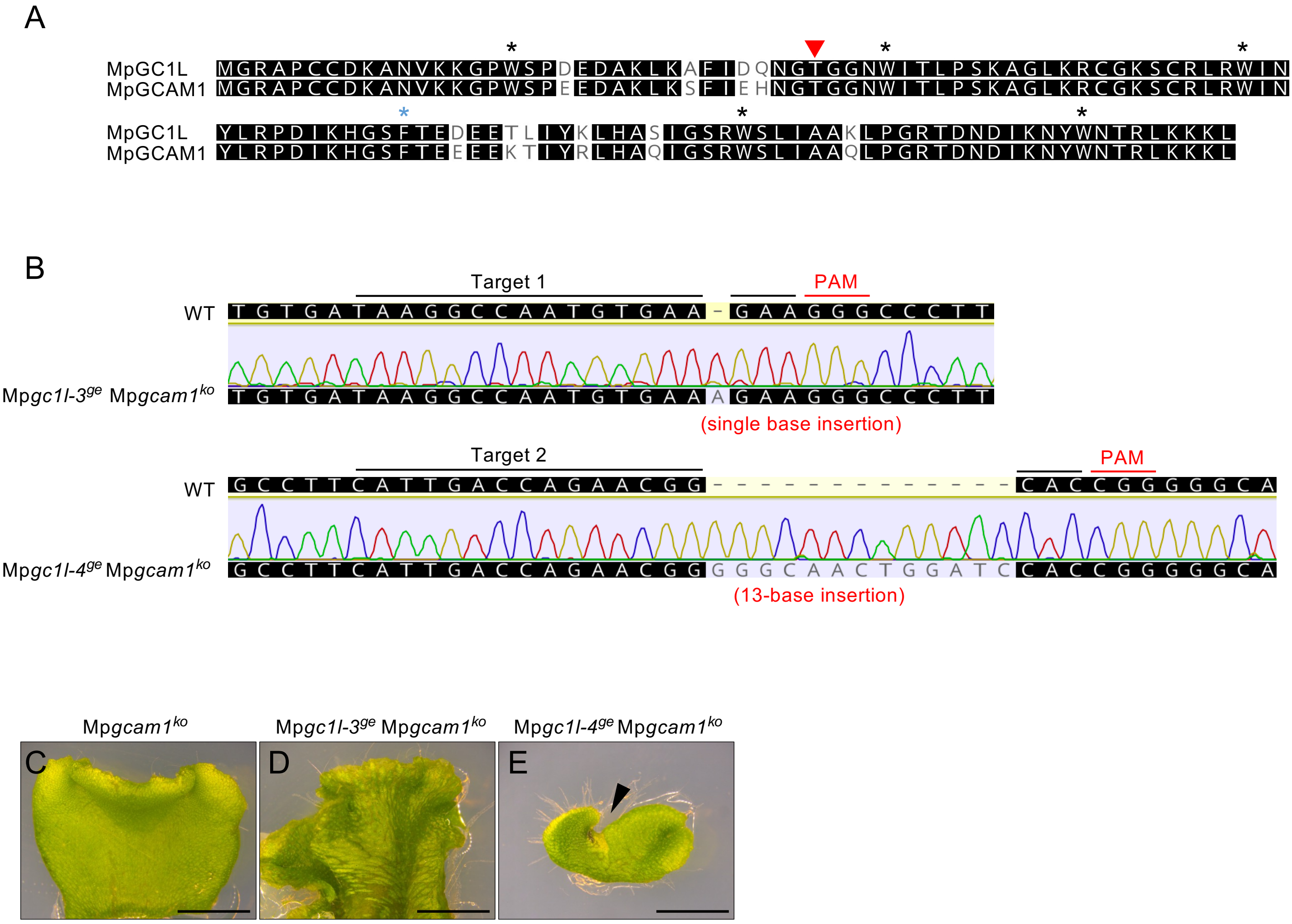


**Figure S4. Sequences of target sites and phenotypes of the Mp*gc1l* Mp*gcam1* double mutants.** (A) Sequence alignment of the R2R3-MYB domains of MpGC1L and MpGCAM1. Black asterisks indicate conserved Trp residues in plant R2R3-MYBs, and blue asterisk indicates a Phe residue conserved in R2R3-MYB subfamily 14. R2R3-MYBs in subfamily 14 are characterized by an additional amino acid insertion between the first and second Trp residues of the R2 repeat compared with other R2R3-MYB subfamilies; the red triangle indicates the position of the insertion. (B) Genome sequences of target sites in the Mp*GC1L* locus in the Mp*gc1l^{ge}* Mp*gcam1^{ko}* mutants. (C-E) Thalli of the Mp*gcam1^{ko}* (C), Mp*gc1l-3^{ge}* Mp*gcam1^{ko}* (D), and Mp*gc1l-4^{ge}* Mp*gcam1^{ko}* mutants (E). An exhausted meristem occasionally observed in Mp*gc1l-4^{ge}* Mp*gcam1^{ko}* is indicated by an arrowhead. Bars = 2 mm.

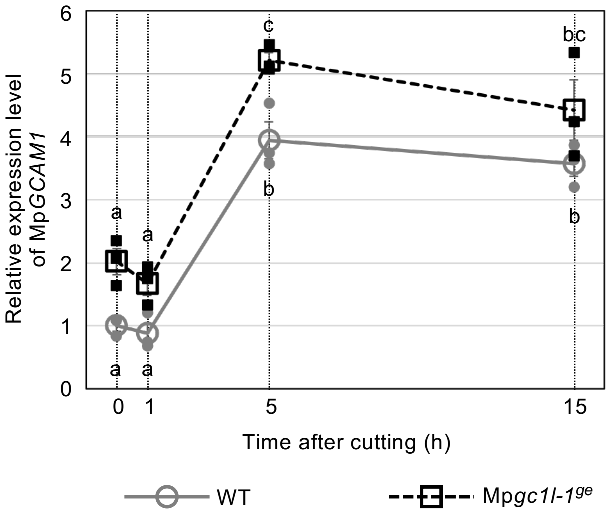


**Figure S5. *GCAM1* expression levels during regeneration.**
RT-qPCR analysis of Mp*GCAM1* expression in gemmalings after cutting off the apical meristem. The apical meristems were removed from 10-d-old WT and Mp*gc1l-1*$^{ge}$ gemmalings, and the basal explants were cultured for the indicated time. Mp*EF1α* was used for normalization. Data are mean ± SE. Different letters above the markers indicate significant differences at $p < 0.05$ by Tukey–Kramer test ($n$ = 3 biological replicates).

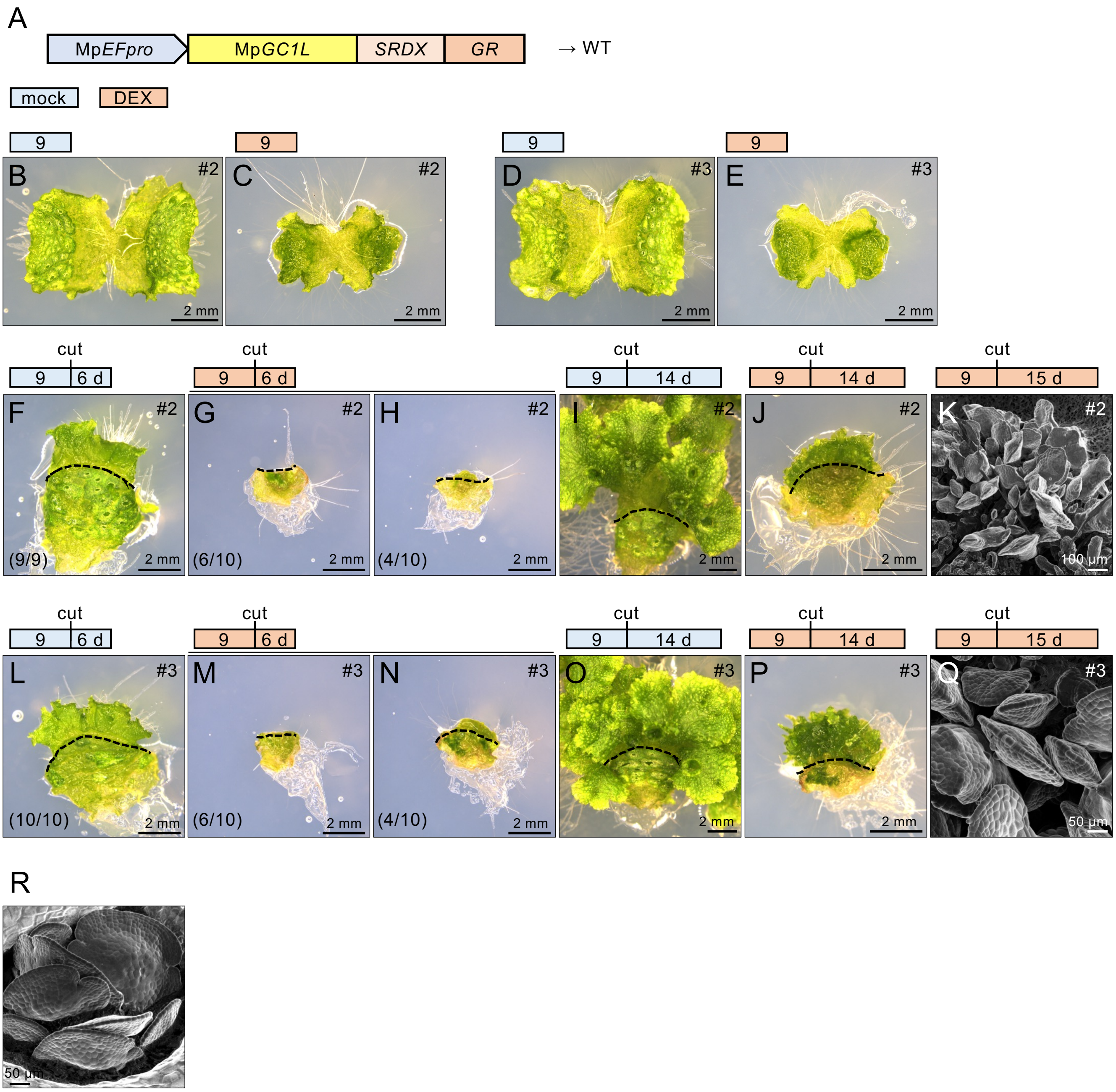


**Figure S6. The effects of introducing Mp*GC1L-SRDX-GR* in WT plants.**
(A) Diagram of the Mp*EFpro:*Mp*GC1L-SRDX-GR* construct, which was introduced into WT plants. (B-E) 9-day-old Mp*EFpro:*Mp*GC1L-SRDX-GR*/WT plants subjected to mock (B and D) or 10 µM DEX treatment (C and E). Mp*EFpro:*Mp*GC1L-SRDX-GR*/WT #2 (B and C) and #3 (D and E). (F-Q) The effects of MpGC1L-SRDX-GR on regeneration in WT plants. The apical meristems were removed from 9-day-old Mp*EFpro:*Mp*GC1L-SRDX-GR*/WT plants subjected to mock (F, I, L, and O) or DEX treatment (G, H, J, K, M, N, P, and Q). The basal explants were subjected to mock (F, I, L and O) or DEX (G, H, J, K, M, N, P, and Q) treatment for 6 d (F-H and L-N), 14 d (I, J, O and P) or 15 d (K and Q). High-magnification images of gemma-like structures were taken by scanning electron microscopy (K and Q). The fraction in brackets shows the number of explants with (F, H, L , and N) or without (G and M) regenerated regions. Mp*EFpro:*Mp*GC1L-SRDX-GR*/WT #2 (F-K) and #3 (L-Q). The bars above the photographs show the DEX treatment conditions. Blue bars, mock treatment; red bars, DEX treatment. The dotted lines shows the border between the basal explant and regenerated region (F-J and L-P). (R) Gemmae developing in a gemma cup in a mock-treated Mp*EFpro:*Mp*GC1L-SRDX-GR*/WT #2 plant taken by scanning electron microscopy.

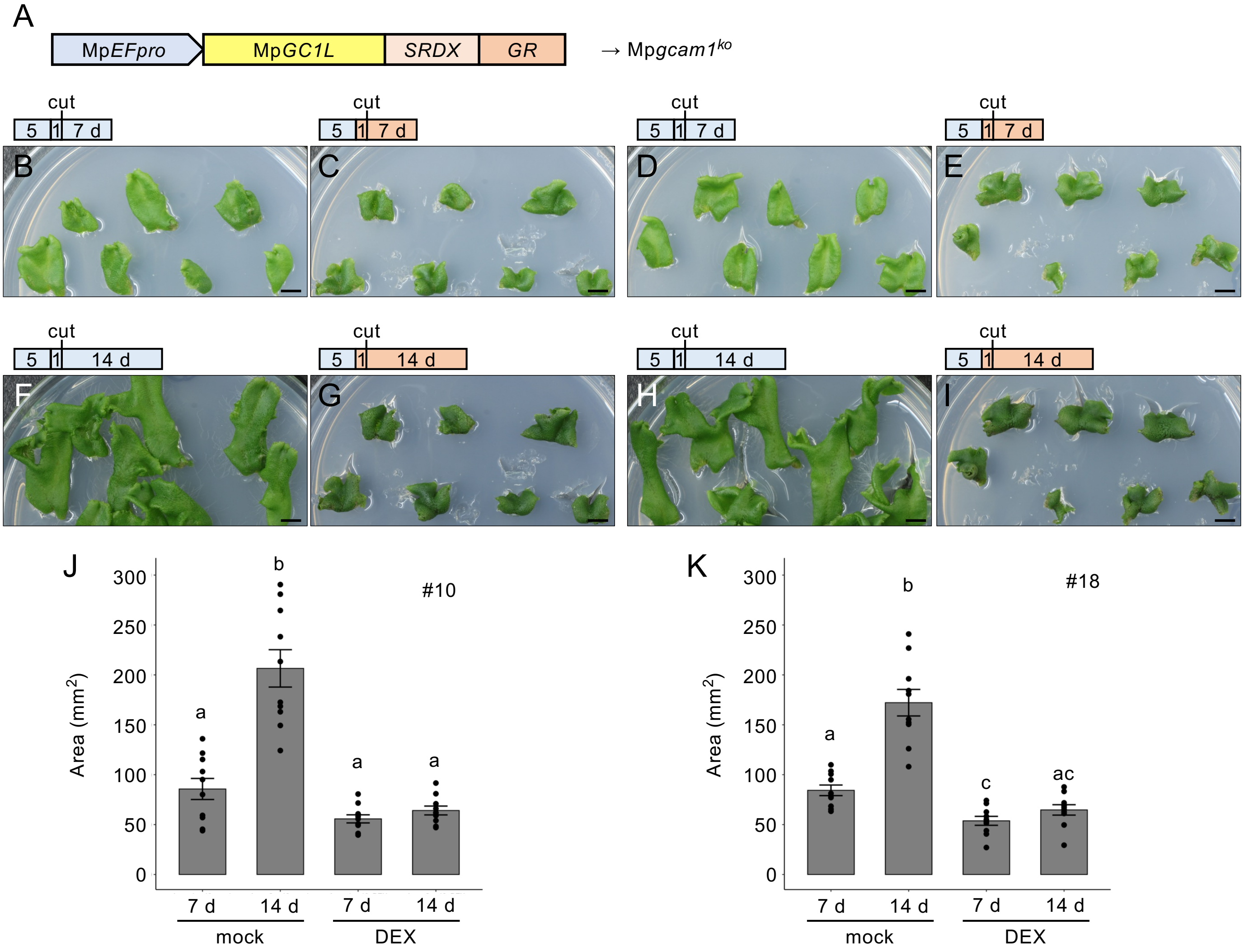


**Figure S7. The effects of introducing Mp*GC1L-SRDX-GR* on apical growth in Mp*gcam1*$^{ko}$ plants.** (A) Schematic representation of the Mp*EFpro*:Mp*GC1L-SRDX-GR* construct. (B-K) The tips of Mp*EFpro*:Mp*GC1L-SRDX-GR* transgenic plants were cultured on control medium for 5 days, transferred to mock (B, D, F and H) or DEX (C, E, G and I) medium, cultured for one day, and the basal explants cut off. The apical explants were subjected to mock (B, D, F and H) or DEX treatment (C, E, G, and I) for 7 d (B-E) or 14 d (F-I). (J and K) Quantification of apical explant area in B-I. Different letters above the bars indicate significant differences (Games-Howell test following Welch's one-way ANOVA, $p < 0.05$). n = 10. Mp*EFpro*:Mp*GC1L-SRDX-GR*/Mp*gcam1*$^{ko}$ #10 (B, C, F, G, and J) and #18 (D, E, H, I, and K).

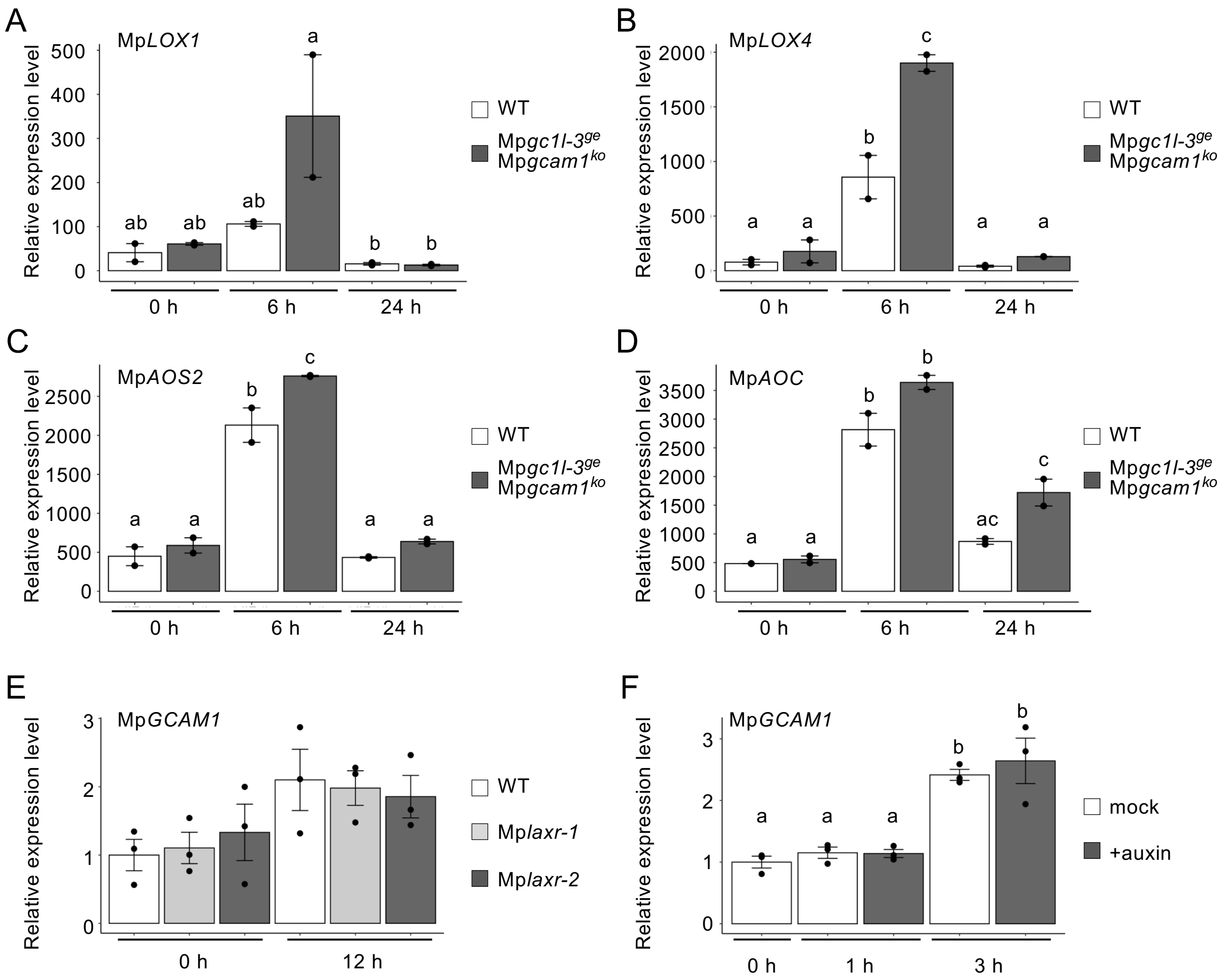


**Figure S8. Expression analysis of genes in known regeneration pathways in basal explants after excision.**
(A-D) Relative expression levels of Mp*LOX1* (A), Mp*LOX4* (B), Mp*AOS2* (C), and Mp*AOC* (D) in WT and Mp*gc1l-3*$^{ge}$ Mp*gcam1*$^{ko}$ based on RNA-seq data. (E and F) Relative expression levels of Mp*GCAM1* determined by RT-qPCR, in the WT and Mp*laxr* mutants (E), and in the WT with or without auxin treatment (F). Mp*EF1α* was used for normalization (E and F). Data are mean ± SE. Different letters above the bars indicate significant differences at $p < 0.05$ by Tukey–Kramer test. Dots indicate biological replicates [$n = 2$ in (*A-D*); $n = 3$ in (E and F)].

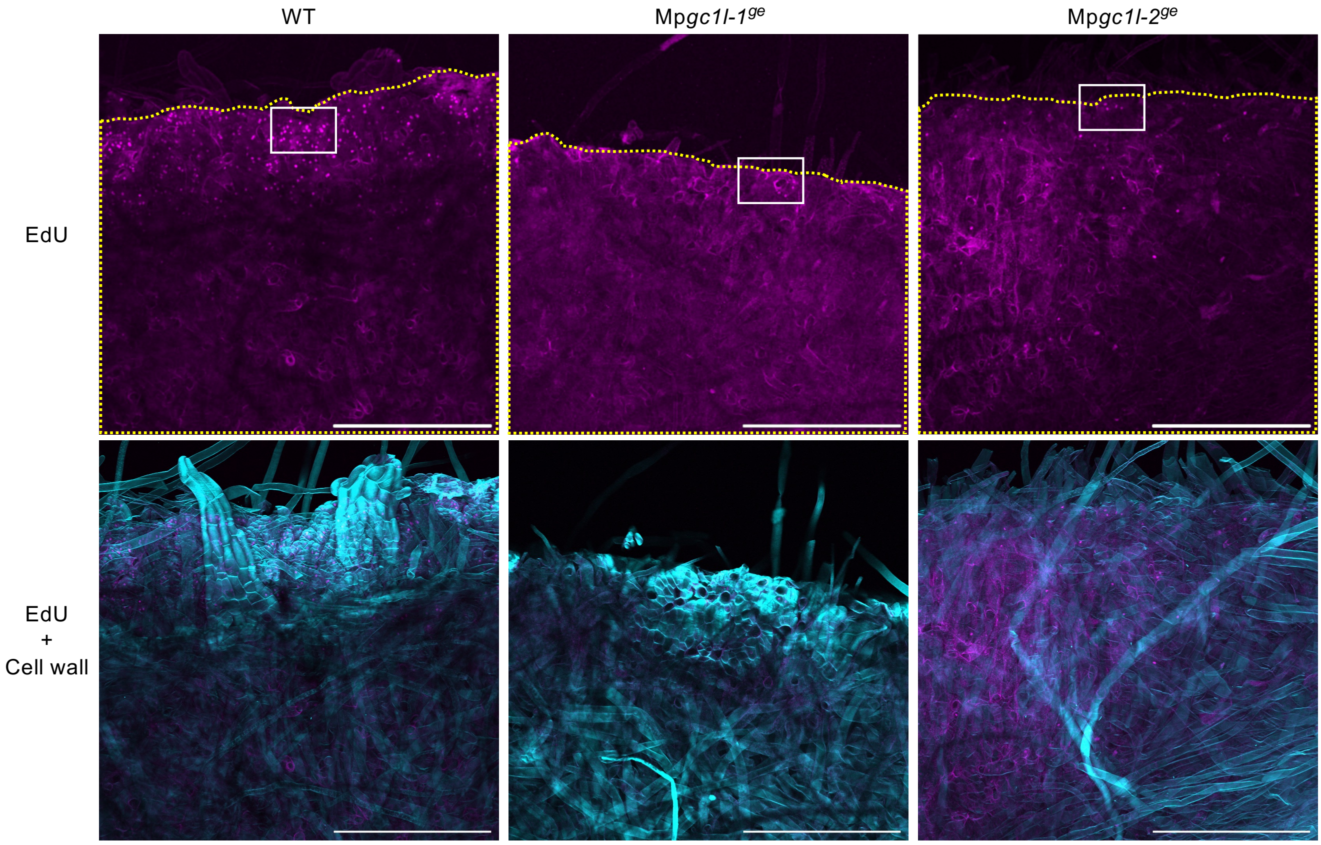


**Figure S9. EdU incorporation assay.**
Representative z-projections of confocal images of basal explants labeled with EdU Alexa Fluor 555 (magenta) and SR2200 (cyan) are shown. Magnified views of the regions marked by white rectangles are shown in Fig. 6*H*. The number of EdU-positive nuclei within the plant area (outlined with yellow dashed lines, $mm^2$) for each line is shown in Fig. 6*I*. Bars = 500 mm.

Table S1. 49 genes down-regulated in the *gc1/gcam1* double mutant at both 6 and 24 hours in cluster_2

| geneID | log2 fold change 6h | log2 fold change 24h | functional annotation | At_best-hit_ID | At_best-hit_symbol | cluster 3 (Ikeuchi et al., 2017) |
|---|---|---|---|---|---|---|
| Mp6g20250 | -6.9346997 | -4.490023 | SUPERFAMILY:SSF53098:Ribonuclease H-like; MapolyID:Mapoly0045s0039 | | | NO |
| Mp2g03770 | -1.9532113 | -3.1967141 | Pfam:PF03732:Retrotransposon gag protein; MapolyID:Mapoly0031s0033 | | | NO |
| Mp3g18310 | -1.4483004 | -1.3248258 | KEGG:K14823:EBP2, EBNA1BP2, rRNA-processing protein EBP2; KOG:KOG3080:Nucleolar protein-like/EBNA1-binding protein, N-term missing, [A]; MobiDBLite:consensus disorder prediction; Coils:Coil; PANTHER:PTHR13028:RRNA PROCESSING PROTEIN EBNA1-BINDING PROTEIN-RELATED; Pfam:PF05890:Eukaryotic rRNA processing protein EBP2; MapolyID:Mapoly0140s0011 | AT3G22660 | EBP2 | YES |
| Mp7g14560 | -1.0952043 | -0.8407021 | PANTHER:PTHR23216:NUCLEOLAR AND COILED-BODY PHOSPHOPROTEIN 1; MobiDBLite:consensus disorder prediction; Pfam:PF05022:SRP40, C-terminal domain; ProSiteProfiles:PS50896:LIS1 homology (LisH) motif profile.; PTHR23216:SF1:NUCLEOLAR AND COILED-BODY PHOSPHOPROTEIN 1; GO:0005515:protein binding; MapolyID:Mapoly0009s0141 | AT5G57120 | nosymbolavailable | YES |
| Mp1g00640 | -1.0020272 | -1.1397963 | KEGG:K00555:TRMT1, trm1, tRNA (guanine26-N2/guanine27-N2)-dimethyltransferase [EC:2.1.1.215 2.1.1.216]; KOG:KOG1253:tRNA methyltransferase, [J]; SUPERFAMILY:SSF53335:S-adenosyl-L-methionine-dependent methyltransferases; TIGRFAM:TIGR00308:TRM1: N2,N2-dimethylguanosine tRNA methyltransferase; MobiDBLite:consensus disorder prediction; PTHR10631:SF12:TRNA (GUANINE(26)-N(2))-DIMETHYLTRANSFERASE 1-RELATED; G3DSA:3.30.56.70; ProSiteProfiles:PS51626:Trm1 methyltransferase domain profile.; Pfam:PF02005:N2,N2-dimethylguanosine tRNA methyltransferase; PANTHER:PTHR10631:N 2 ,N 2 -DIMETHYLGUANOSINE TRNA METHYLTRANSFERASE; CDD:cd02440:AdoMet_MTases; GO:0003723:RNA binding; GO:0008033:tRNA processing; GO:0004809:tRNA (guanine-N2-)-methyltransferase activity; MapolyID:Mapoly0103s0023 | AT3G02320 | nosymbolavailable | YES |
| Mp5g11380 | -0.9491499 | -1.6514257 | MobiDBLite:consensus disorder prediction; PANTHER:PTHR33227; Pfam:PF04885:Stigma-specific protein, Stig1; MapolyID:Mapoly0093s0061 | AT5G55110 | nosymbolavailable | NO |
| Mp4g08260 | -0.9212097 | -0.8438149 | KOG:KOG0339:ATP-dependent RNA helicase, [A]; G3DSA:3.40.50.300; Pfam:PF00271:Helicase conserved C-terminal domain; ProSiteProfiles:PS51194:Superfamilies 1 and 2 helicase C-terminal domain profile.; PTHR47958:SF8:ATP-DEPENDENT RNA HELICASE DBP3 ISOFORM X1; ProSiteProfiles:PS51192:Superfamilies 1 and 2 helicase ATP-binding type-1 domain profile.; PANTHER:PTHR47958:ATP-DEPENDENT RNA HELICASE DBP3; MobiDBLite:consensus disorder prediction; Pfam:PF00270:DEAD/DEAH box helicase; SUPERFAMILY:SSF52540:P-loop containing nucleoside triphosphate hydrolases; GO:0003676:nucleic acid binding; GO:0005524:ATP binding; MapolyID:Mapoly0120s0020 | AT2G28600 | nosymbolavailable | YES |
| Mp5g01410 | -0.9020191 | -1.0474717 | KEGG:K01859:E5.5.1.6, chalcone isomerase [EC:5.5.1.6]; G3DSA:3.50.70.10; G3DSA:1.10.890.20; SUPERFAMILY:SSF54626:Chalcone isomerase; Pfam:PF02431:Chalcone-flavanone isomerase; PANTHER:PTHR47588:CHALCONE--FLAVONONE ISOMERASE 3-RELATED; GO:0016872:intramolecular lyase activity; MapolyID:Mapoly0175s0004 | AT5G05270 | CHIL,AtCHIL | NO |
| Mp5g08210 | -0.86537 | -0.6863308 | KEGG:K11128:GAR1, NOLA1, H/ACA ribonucleoprotein complex subunit 1; KOG:KOG3262:H/ACA small nucleolar RNP component GAR1, C-term missing, [J]; Pfam:PF04410:Gar1/Naf1 RNA binding region; MobiDBLite:consensus disorder prediction; SUPERFAMILY:SSF50447:Translation proteins; PANTHER:PTHR23237:NUCLEOLAR PROTEIN FAMILY A MEMBER 1 SNORNP PROTEIN GAR1; G3DSA:2.40.10.230:Probable tRNA pseudouridine synthase domain; PTHR23237:SF12:H/ACA RIBONUCLEOPROTEIN COMPLEX SUBUNIT; GO:0042254:ribosome biogenesis; GO:0001522:pseudouridine synthesis; MapolyID:Mapoly0086s0024 | AT3G03920 | nosymbolavailable | YES |
| Mp1g29710 | -0.8533166 | -0.9574988 | MobiDBLite:consensus disorder prediction; G3DSA:3.10.450.40; Pfam:PF11523:Protein of unknown function (DUF3223); PANTHER:PTHR33415; PTHR33415:SF12:PROTEIN EMBRYO DEFECTIVE 514; MapolyID:Mapoly0139s0003 | AT5G62440 | nosymbolavailable | YES |
| Mp4g18120 | -0.8476624 | -0.6814108 | KEGG:K14767:UTP3, SAS10, U3 small nucleolar RNA-associated protein 3; KOG:KOG3117:Protein involved in rRNA processing, [A]; MobiDBLite:consensus disorder prediction; Coils:Coil; PTHR13237:SF8:SOMETHING ABOUT SILENCING PROTEIN 10; PANTHER:PTHR13237:SOMETHING ABOUT SILENCING PROTEIN 10-RELATED; Pfam:PF04000:Sas10/Utp3/C1D family; Pfam:PF09368:Sas10 C-terminal domain; MapolyID:Mapoly0041s0093 | AT2G43650 | EMB2777 | YES |
| Mp4g17130 | -0.8330922 | -0.9547095 | KOG:KOG2974:Uncharacterized conserved protein, N-term missing, [S]; MobiDBLite:consensus disorder prediction; Coils:Coil; PANTHER:PTHR13245:RRP15-LIKE PROTEIN; Pfam:PF07890:Rrp15p; GO:0006364:rRNA processing; MapolyID:Mapoly0148s0006 | AT5G48240 | nosymbolavailable | NO |
| Mp8g07360 | -0.8286827 | -0.6845574 | KEGG:K11129:NHP2, NOLA2, H/ACA ribonucleoprotein complex subunit 2; KOG:KOG3167:Box H/ACA snoRNP component, involved in ribosomal RNA pseudouridinylation, [A]; PANTHER:PTHR23105:RIBOSOMAL PROTEIN L7AE FAMILY MEMBER; PRINTS:PR00883:High mobility group-like nuclear protein signature; G3DSA:3.30.1330.30; SUPERFAMILY:SSF55315:L30e-like; ProSitePatterns:PS01082:Ribosomal protein L7Ae signature.; Pfam:PF01248:Ribosomal protein L7Ae/L30e/S12e/Gadd45 family; PRINTS:PR00881:Ribosomal protein L7A/RS6 family signature; PTHR23105:SF146; GO:0003723:RNA binding; GO:1990904:ribonucleoprotein complex; GO:0042254:ribosome biogenesis; GO:0005730:nucleolus; MapolyID:Mapoly0013s0057 | AT5G08180 | nosymbolavailable | YES |
| Mp1g07840 | -0.8223603 | -0.737731 | KEGG:K14852:RRS1, regulator of ribosome biosynthesis; KOG:KOG1765:Regulator of ribosome synthesis, [J]; PANTHER:PTHR17602:RIBOSOME BIOGENESIS REGULATORY PROTEIN; MobiDBLite:consensus disorder prediction; Pfam:PF04939:Ribosome biogenesis regulatory protein (RRS1); PTHR17602:SF5:RIBOSOME BIOGENESIS REGULATORY PROTEIN; Coils:Coil; GO:0005634:nucleus; GO:0042254:ribosome biogenesis; MapolyID:Mapoly0036s0028 | AT2G37990 | nosymbolavailable | YES |
| Mp6g04100 | -0.8142416 | -0.765291 | KEGG:K14835:NOP2, 25S rRNA (cytosine2870-C5)-methyltransferase [EC:2.1.1.310]; KOG:KOG1122:tRNA and rRNA cytosine-C5-methylase (nucleolar protein NOL1/NOP2), [A]; Pfam:PF01189:16S rRNA methyltransferase RsmB/F; G3DSA:3.30.70.3130; MobiDBLite:consensus disorder prediction; Pfam:PF17125:N-terminal domain of 16S rRNA methyltransferase RsmF; G3DSA:3.40.50.150:Vaccinia Virus protein VP39; PANTHER:PTHR22807:NOP2 YEAST -RELATED NOL1/NOP2/FMU SUN DOMAIN-CONTAINING; ProSiteProfiles:PS51686:SAM-dependent MTase RsmB/NOP-type domain profile.; SUPERFAMILY:SSF53335:S-adenosyl-L-methionine-dependent methyltransferases; TIGRFAM:TIGR00446:nop2p: NOL1/NOP2/sun family putative RNA methylase; PTHR22807:SF65:BNACNNG49010D PROTEIN; PRINTS:PR02008:RNA (C5-cytosine) methyltransferase signature; PRINTS:PR02012:RNA (C5-cytosine) methyltransferase NOP2 subfamily signature; GO:0008757:S-adenosylmethionine-dependent methyltransferase activity; GO:0008168:methyltransferase activity; GO:0003723:RNA binding; GO:0006396:RNA processing; GO:0001510:RNA methylation; MapolyID:Mapoly0034s0108 | AT4G26600 | NOP2B | YES |
| Mp8g08350 | -0.7548541 | -0.7349539 | KEGG:K07565:NIP7, 60S ribosome subunit biogenesis protein NIP7; KOG:KOG3492:Ribosome biogenesis protein NIP7, [J]; G3DSA:3.10.450.220; SUPERFAMILY:SSF88802:Pre-PUA domain; Pfam:PF17833:UPF0113 Pre-PUA domain; Pfam:PF03657:UPF0113 PUA domain; ProSiteProfiles:PS50890:PUA domain profile.; PTHR23415:SF4:60S RIBOSOME SUBUNIT BIOGENESIS PROTEIN NIP7 HOMOLOG; PIRSF:PIRSF017190:NIP7; PANTHER:PTHR23415:CYCLIN-DEPENDENT KINASES REGULATORY SUBUNIT/60S RIBOSOME SUBUNIT BIOGENESIS PROTEIN NIP7; SUPERFAMILY:SSF88697:PUA domain-like; SMART:SM00359:pua_5; G3DSA:2.30.130.10; GO:0005634:nucleus; GO:0003723:RNA binding; GO:0042255:ribosome assembly; MapolyID:Mapoly0063s0083 | AT4G15770 | nosymbolavailable | YES |
| Mp3g19910 | -0.7453449 | -0.5505229 | MobiDBLite:consensus disorder prediction; Coils:Coil; Pfam:PF10441:Urb2/Npa2 family; PANTHER:PTHR15682:UNHEALTHY RIBOSOME BIOGENESIS PROTEIN 2 HOMOLOG; MapolyID:Mapoly0049s0043 | AT4G30150 | nosymbolavailable | YES |
| Mp7g06450 | -0.7317512 | -0.719444 | KEGG:K06173:truA, PUS1, tRNA pseudouridine38-40 synthase [EC:5.4.99.12]; KOG:KOG2553:Pseudouridylate synthase, [J]; PANTHER:PTHR11142:PSEUDOURIDYLATE SYNTHASE; SUPERFAMILY:SSF55120:Pseudouridine synthase; G3DSA:3.30.70.580; Coils:Coil; G3DSA:3.30.70.660; PTHR11142:SF4:TRNA PSEUDOURIDINE SYNTHASE A; MobiDBLite:consensus disorder prediction; Pfam:PF01416:tRNA pseudouridine synthase; CDD:cd02568:PseudoU_synth_PUS1_PUS2; GO:0031119:tRNA pseudouridine synthesis; GO:0009982:pseudouridine synthase activity; GO:0001522:pseudouridine synthesis; GO:0003723:RNA binding; GO:0009451:RNA modification; MapolyID:Mapoly0057s0025 | AT1G20370 | nosymbolavailable | YES |
| Mp6g02290 | -0.731611 | -1.2130028 | KEGG:K00134:GAPDH, gapA, glyceraldehyde 3-phosphate dehydrogenase [EC:1.2.1.12]; KOG:KOG0657:Glyceraldehyde 3-phosphate dehydrogenase, [G]; PANTHER:PTHR10836:GLYCERALDEHYDE 3-PHOSPHATE DEHYDROGENASE; SUPERFAMILY:SSF55347:Glyceraldehyde-3-phosphate dehydrogenase-like, C-terminal domain; SMART:SM00846:gp_dh_n_7; PRINTS:PR00078:Glyceraldehyde-3-phosphate dehydrogenase signature; PTHR10836:SF76:GLYCERALDEHYDE-3-PHOSPHATE DEHYDROGENASE-RELATED; Pfam:PF00044:Glyceraldehyde 3-phosphate dehydrogenase, NAD binding domain; G3DSA:3.40.50.720; G3DSA:3.30.360.10:Dihydrodipicolinate Reductase, domain 2; SUPERFAMILY:SSF51735:NAD(P)-binding Rossmann-fold domains; ProSitePatterns:PS00071:Glyceraldehyde 3-phosphate dehydrogenase active site.; Pfam:PF02800:Glyceraldehyde 3-phosphate dehydrogenase, C-terminal domain; TIGRFAM:TIGR01534:GAPDH-I: glyceraldehyde-3-phosphate dehydrogenase, type I; GO:0016620:oxidoreductase activity, acting on the aldehyde or oxo group of donors, NAD or NADP as acceptor; GO:0006006:glucose metabolic process; GO:0051287:NAD binding; GO:0050661:NADP binding; MapolyID:Mapoly0035s0014 | AT1G16300 | GAPCP-2 | NO |

| | | | | | | |
|---|---|---|---|---|---|---|
| Mp4g00830 | -0.7313131 | -0.8951967 | KEGG:K14549:UTP15, U3 small nucleolar RNA-associated protein 15; KOG:KOG0310:Conserved WD40 repeat-containing protein, [S]; G3DSA:2.130.10.10; PANTHER:PTHR19924:UTP15 U3 SMALL NUCLEOLAR RNA-ASSOCIATED PROTEIN 15 FAMILY MEMBER; SMART:SM00320:WD40_4; Pfam:PF00400:WD domain, G-beta repeat; ProSiteProfiles:PS50082:Trp-Asp (WD) repeats profile.; ProSiteProfiles:PS50294:Trp-Asp (WD) repeats circular profile.; SUPERFAMILY:SSF50978:WD40 repeat-like; CDD:cd00200:WD40; Pfam:PF09384:UTP15 C terminal; PTHR19924:SF26:U3 SMALL NUCLEOLAR RNA-ASSOCIATED PROTEIN 15 HOMOLOG; GO:0005730:nucleolus; GO:0005515:protein binding; GO:0006364:rRNA processing; MapolyID:Mapoly0066s0060 | AT2G47990 | SWA1,EDA13,EDA19 | YES |
| Mp5g18010 | -0.7159767 | -0.9651964 | KEGG:K14826:FPR3_4, FK506-binding nuclear protein [EC:5.2.1.8]; KOG:KOG0544:FKBP-type peptidyl-prolyl cis-trans isomerase, [O]; MobiDBLite:consensus disorder prediction; G3DSA:3.10.50.40; SUPERFAMILY:SSF54534:FKBP-like; SUPERFAMILY:SSF69203:Nucleoplasmin-like core domain; G3DSA:2.60.120.340; PTHR43811:SF47:PEPTIDYLPROLYL ISOMERASE; PANTHER:PTHR43811:FKBP-TYPE PEPTIDYL-PROLYL CIS-TRANS ISOMERASE FKPA; ProSiteProfiles:PS50059:FKBP-type peptidyl-prolyl cis-trans isomerase domain profile.; Pfam:PF17800:Nucleoplasmin-like domain; Pfam:PF00254:FKBP-type peptidyl-prolyl cis-trans isomerase; PIRSF:PIRSF001473:FK506-bp_FPR3; GO:0003755:peptidyl-prolyl cis-trans isomerase activity; MapolyID:Mapoly0084s0048 | AT4G25340 | ATFKBP53,FKBP53 | YES |
| Mp2g02550 | -0.7120231 | -0.9724158 | KEGG:K02260:COX17, cytochrome c oxidase assembly protein subunit 17; KOG:KOG3496:Cytochrome c oxidase assembly protein/Cu2+ chaperone COX17, N-term missing, [O]; SUPERFAMILY:SSF47072:Cysteine alpha-hairpin motif; PANTHER:PTHR16719:CYTOCHROME C OXIDASE COPPER CHAPERONE; MobiDBLite:consensus disorder prediction; PTHR16719:SF0:CYTOCHROME C OXIDASE COPPER CHAPERONE; Pfam:PF05051:Cytochrome C oxidase copper chaperone (COX17); G3DSA:1.10.287.1130:CytochromE C oxidase copper chaperone; ProSiteProfiles:PS51808:Coiled coil-helix-coiled coil-helix (CHCH) domain profile.; GO:0005507:copper ion binding; GO:0005758:mitochondrial intermembrane space; GO:0016531:copper chaperone activity; MapolyID:Mapoly0075s0017 | AT1G53030 | nosymbolavailable | NO |
| Mp3g15820 | -0.7118 | -0.7119986 | KOG:KOG0216:RNA polymerase I, second largest subunit, [K]; G3DSA:2.40.50.150; Pfam:PF04563:RNA polymerase beta subunit; PANTHER:PTHR20856:DNA-DIRECTED RNA POLYMERASE I SUBUNIT 2; ProSitePatterns:PS01166:RNA polymerases beta chain signature.; G3DSA:3.90.1070.20; PTHR20856:SF5:DNA-DIRECTED RNA POLYMERASE I SUBUNIT RPA2; G3DSA:3.90.1110.10; G3DSA:3.90.1100.10; Pfam:PF04561:RNA polymerase Rpb2, domain 2; Pfam:PF04565:RNA polymerase Rpb2, domain 3; G3DSA:3.90.1800.10:RNA polymerase alpha subunit dimerisation domain; CDD:cd00653:RNA_pol_B_RPB2; SUPERFAMILY:SSF64484:beta and beta-prime subunits of DNA dependent RNA-polymerase; G3DSA:2.40.270.10; Pfam:PF06883:RNA polymerase I, Rpa2 specific domain; Pfam:PF04560:RNA polymerase Rpb2, domain 7; Pfam:PF00562:RNA polymerase Rpb2, domain 6; GO:0006351:transcription, DNA-templated; GO:0005634:nucleus; GO:0003677:DNA binding; GO:0003899:DNA-directed 5'-3' RNA polymerase activity; GO:0032549:ribonucleoside binding; MapolyID:Mapoly0004s0090 | AT1G29940 | NRPA2 | YES |
| Mp7g01710 | -0.7039101 | -0.6234427 | KEGG:K14844:PUF6, pumilio homology domain family member 6; KOG:KOG2050:Puf family RNA-binding protein, [J]; MobiDBLite:consensus disorder prediction; ProSiteProfiles:PS50302:Pumilio RNA-binding repeat profile.; Pfam:PF00806:Pumilio-family RNA binding repeat; G3DSA:1.25.10.10; ProSiteProfiles:PS50303:Pumilio homology domain (PUM-HD) profile.; PANTHER:PTHR13389:UNCHARACTERIZED; SUPERFAMILY:SSF48371:ARM repeat; SMART:SM00025:pum_5; Pfam:PF08144:CPL (NUC119) domain; GO:0003723:RNA binding; MapolyID:Mapoly0099s0044 | AT3G16810 | APUM24,PUM24 | YES |
| Mp1g04820 | -0.6991359 | -0.698055 | KEGG:K11090:LA, SSB, lupus La protein; KOG:KOG1855:Predicted RNA-binding protein, N-term missing, C-term missing, [R]; PTHR22792:SF79:OS02G0610400 PROTEIN; MobiDBLite:consensus disorder prediction; Pfam:PF08777:RNA binding motif; ProSiteProfiles:PS50102:Eukaryotic RNA Recognition Motif (RRM) profile.; Pfam:PF05383:La domain; SUPERFAMILY:SSF54928:RNA-binding domain, RBD; CDD:cd12291:RRM1_La; Pfam:PF00076:RNA recognition motif. (a.k.a. RRM, RBD, or RNP domain); SMART:SM00715:la; G3DSA:1.10.10.10:""winged helix"" repressor DNA binding domain; SMART:SM00360:rrm1_1; G3DSA:3.30.70.330; PANTHER:PTHR22792:LUPUS LA PROTEIN-RELATED; ProSiteProfiles:PS50961:La-type HTH domain profile.; SUPERFAMILY:SSF46785:"Winged helix" DNA-binding domain; CDD:cd08030:LA_like_plant; PRINTS:PR00302:Lupus La protein signature; GO:0005634:nucleus; GO:0003723:RNA binding; GO:0003676:nucleic acid binding; GO:1990904:ribonucleoprotein complex; GO:0006396:RNA processing; MapolyID:Mapoly0005s0125 | AT4G32720 | AtLa1,La1 | YES |
| Mp7g18590 | -0.69505 | -0.5159525 | KEGG:K14808:DDX54, DBP10, ATP-dependent RNA helicase DDX54/DBP10 [EC:3.6.4.13]; KOG:KOG0337:ATP-dependent RNA helicase, C-term missing, [A]; ProSiteProfiles:PS51195:DEAD-box RNA helicase Q motif profile.; ProSiteProfiles:PS51192:Superfamilies 1 and 2 helicase ATP-binding type-1 domain profile.; Pfam:PF08147:DBP10CT (NUC160) domain; G3DSA:3.40.50.300; Coils:Coil; MobiDBLite:consensus disorder prediction; ProSiteProfiles:PS51194:Superfamilies 1 and 2 helicase C-terminal domain profile.; SMART:SM00490:helicmild6; SUPERFAMILY:SSF52540:P-loop containing nucleoside triphosphate hydrolases; Pfam:PF00270:DEAD/DEAH box helicase; PANTHER:PTHR47959:ATP-DEPENDENT RNA HELICASE RHLE-RELATED; CDD:cd18787:SF2_C_DEAD; Pfam:PF00271:Helicase conserved C-terminal domain; ProSitePatterns:PS00039:DEAD-box subfamily ATP-dependent helicases signature.; SMART:SM00487:ultradead3; CDD:cd17959:DEADc_DDX54; PTHR47959:SF8:DEAD-BOX ATP-DEPENDENT RNA HELICASE 29; SMART:SM01123:DBP10CT_2; GO:0004386:helicase activity; GO:0005634:nucleus; GO:0003723:RNA binding; GO:0003676:nucleic acid binding; GO:0003724:RNA helicase activity; GO:0005524:ATP binding; MapolyID:Mapoly0165s0019 | AT1G77030 | nosymbolavailable | YES |
| Mp8g15160 | -0.6907251 | -0.6165437 | KEGG:K11108:RCL1, RNA 3'-terminal phosphate cyclase-like protein; KOG:KOG3980:RNA 3'-terminal phosphate cyclase, [A]; Pfam:PF05189:RNA 3'-terminal phosphate cyclase (RTC), insert domain; PANTHER:PTHR11096:RNA 3' TERMINAL PHOSPHATE CYCLASE; ProSitePatterns:PS01287:RNA 3'-terminal phosphate cyclase signature.; CDD:cd00875:RNA_Cyclase_Class_I; SUPERFAMILY:SSF55205:EPT/RTPC-like; G3DSA:3.30.360.20; PIRSF:PIRSF005378:RNA_3-term_P_cyclase; Pfam:PF01137:RNA 3'-terminal phosphate cyclase; G3DSA:3.65.10.20; PTHR11096:SF1:RNA 3'-TERMINAL PHOSPHATE CYCLASE-LIKE PROTEIN; TIGRFAM:TIGR03400:18S_RNA_Rcl1p: 18S rRNA biogenesis protein RCL1; GO:0005730:nucleolus; GO:0042254:ribosome biogenesis; GO:0006396:RNA processing; GO:0003824:catalytic activity; MapolyID:Mapoly0187s0002 | AT5G22100 | nosymbolavailable | YES |
| Mp5g13270 | -0.6839842 | -0.5840674 | KEGG:K00602:purH, phosphoribosylaminoimidazolecarboxamide formyltransferase / IMP cyclohydrolase [EC:2.1.2.3 3.5.4.10]; KOG:KOG2555:AICAR transformylase/IMP cyclohydrolase/methylglyoxal synthase, [F]; PANTHER:PTHR11692:BIFUNCTIONAL PURINE BIOSYNTHESIS PROTEIN PURH; SMART:SM00798:aicarft_impchas; CDD:cd01421:IMPCH; SMART:SM00851:MGS_2a; Pfam:PF02142:MGS-like domain; G3DSA:3.40.140.20; TIGRFAM:TIGR00355:purH: phosphoribosylaminoimidazolecarboxamide formyltransferase/IMP cyclohydrolase; Pfam:PF01808:AICARFT/IMPCHase bienzyme; Hamap:MF_00139:Bifunctional purine biosynthesis protein PurH [purH].; ProSiteProfiles:PS51855:MGS-like domain profile.; G3DSA:3.40.50.1380; PTHR11692:SF1:AICARFT/IMPCHASE BIENZYME FAMILY PROTEIN; SUPERFAMILY:SSF53927:Cytidine deaminase-like; PIRSF:PIRSF000414:PurH; SUPERFAMILY:SSF52335:Methylglyoxal synthase-like; GO:0006164:purine nucleotide biosynthetic process; GO:0003824:catalytic activity; GO:0003937:IMP cyclohydrolase activity; GO:0004643:phosphoribosylaminoimidazolecarboxamide formyltransferase activity; MapolyID:Mapoly0032s0021 | AT2G35040 | nosymbolavailable | YES |
| Mp3g12300 | -0.68347 | -1.2871549 | PANTHER:PTHR31867:EXPANSIN-A15; SUPERFAMILY:SSF50685:Barwin-like endoglucanases; SMART:SM00837:dpbb_1; ProSiteProfiles:PS50842:Expansin, family-45 endoglucanase-like domain profile.; Pfam:PF03330:Lytic transglycolase; PRINTS:PR01225:Expansin/Lol pI family signature; G3DSA:2.40.40.10; ProSiteProfiles:PS50843:Expansin, Cellulose-binding-like domain profile.; PRINTS:PR01226:Expansin signature; G3DSA:2.60.40.760; SUPERFAMILY:SSF49590:PHL pollen allergen; Pfam:PF01357:Expansin C-terminal domain; GO:0005576:extracellular region; GO:0009664:plant-type cell wall organization; MapolyID:Mapoly0050s0035 | AT2G03090 | EXPA15,ATEXP15,ATHEXPALPHA1.3,ATEXPA15,EXP15 | NO |
| Mp5g10240 | -0.6583862 | -0.9133718 | MobiDBLite:consensus disorder prediction; Pfam:PF17800:Nucleoplasmin-like domain; G3DSA:2.60.120.340; ProSitePatterns:PS00028:Zinc finger C2H2 type domain signature.; PANTHER:PTHR31802:32 KDA HEAT SHOCK PROTEIN-RELATED; ProSiteProfiles:PS50157:Zinc finger C2H2 type domain profile.; MapolyID:Mapoly0048s0048 | AT5G22650 | HD2,HDT02,HDA4,HD2B,ATHD2B,HDT2,ATHD2 | YES |
| Mp1g21390 | -0.6576287 | -0.7014245 | KEGG:K11294:NCL, NSR1, nucleolin; KOG:KOG4210:Nuclear localization sequence binding protein, N-term missing, [K]; MobiDBLite:consensus disorder prediction; PTHR24012:SF717:POLYNUCLEOTIDE ADENYLYLTRANSFERASE DOMAIN/RNA RECOGNITION MOTIF PROTEIN-RELATED; PANTHER:PTHR24012:RNA BINDING PROTEIN; CDD:cd12451:RRM2_NUCLs; G3DSA:3.30.70.330; SMART:SM00360:rrm1_1; ProSiteProfiles:PS50102:Eukaryotic RNA Recognition Motif (RRM) profile.; SUPERFAMILY:SSF54928:RNA-binding domain, RBD; Pfam:PF00076:RNA recognition motif. (a.k.a. RRM, RBD, or RNP domain); GO:0003676:nucleic acid binding; MapolyID:Mapoly0001s0474 | AT3G18610 | PARLL1,NUC2,ATNUC-L2,NUC-L2 | YES |
| Mp2g18510 | -0.6547612 | -0.9852464 | Pfam:PF03140:Plant protein of unknown function; PANTHER:PTHR31549:PROTEIN, PUTATIVE (DUF247)-RELATED-RELATED; MapolyID:Mapoly0137s0030 | AT3G50120 | nosymbolavailable | NO |
| Mp3g16040 | -0.6463966 | -0.7048752 | KEGG:K07556:ATPeAF2, ATPAF2, ATP12, ATP synthase mitochondrial F1 complex assembly factor 2; KOG:KOG3015:F1-ATP synthase assembly protein, [C]; PANTHER:PTHR21013:ATP SYNTHASE MITOCHONDRIAL F1 COMPLEX ASSEMBLY FACTOR 2/ATP12 PROTEIN, MITOCHONDRIAL PRECURSOR; SUPERFAMILY:SSF160909:ATP12-like; G3DSA:1.10.3580.10:ATP12 ATPase; Pfam:PF07542:ATP12 chaperone protein; G3DSA:3.30.2180.30; GO:0043461:proton-transporting ATP synthase complex assembly; MapolyID:Mapoly0004s0068 | AT5G40660 | nosymbolavailable | YES |

| | | | | | | |
|---|---|---|---|---|---|---|
| Mp8g04700 | -0.6431153 | -0.6557514 | KOG:KOG1263:Multicopper oxidases, [Q]; CDD:cd13868:CuRO_2_CotA_like; CDD:cd13844:CuRO_1_BOD_CotA_like; SUPERFAMILY:SSF49503:Cupredoxins; PANTHER:PTHR11709:MULTI-COPPER OXIDASE; Pfam:PF07731:Multicopper oxidase; CDD:cd13891:CuRO_3_CotA_like; Pfam:PF07732:Multicopper oxidase; PTHR11709:SF340:MULTICOPPER OXIDASE LPR1 HOMOLOG 1; G3DSA:2.60.40.420; GO:0005507:copper ion binding; GO:0016491:oxidoreductase activity; MapolyID:Mapoly0186s0019 | AT1G23010 | LPR1 | NO |
| Mp6g08990 | -0.6408097 | -0.5286213 | KEGG:K14544:UTP22, NOL6, U3 small nucleolar RNA-associated protein 22; KOG:KOG2054:Nucleolar RNA-associated protein (NRAP), [S]; Pfam:PF17406:Nrap protein PAP/OAS1-like domain 5; Pfam:PF17403:Nrap protein PAP/OAS-like domain; G3DSA:3.30.460.10:Beta Polymerase; Pfam:PF17404:Nrap protein domain 3; PANTHER:PTHR17972:NUCLEOLAR RNA-ASSOCIATED PROTEIN; Pfam:PF03813:Nrap protein domain 1; G3DSA:1.10.1410.10; Pfam:PF17407:Nrap protein domain 6; Pfam:PF17405:Nrap protein nucleotidyltransferase domain 4; MapolyID:Mapoly0060s0020 | AT1G63810 | nosymbolavailable | YES |
| Mp2g12880 | -0.6391241 | -0.8564327 | KEGG:K14563:NOP1, FBL, rRNA 2'-O-methyltransferase fibrillarin [EC:2.1.1.-]; KOG:KOG1596:Fibrillarin and related nucleolar RNA-binding proteins, N-term missing, [A]; PANTHER:PTHR10335:RRNA 2-O-METHYLTRANSFERASE FIBRILLARIN; PIRSF:PIRSF006540:Nop17p; PTHR10335:SF22:FIBRILLARIN, S-ADENOSYL-L-METHIONINE-DEPENDENT METHYLTRANSFERASE-RELATED; Hamap:MF_00351:Fibrillarin-like rRNA/tRNA 2'-O-methyltransferase [flpA].; SUPERFAMILY:SSF53335:S-adenosyl-L-methionine-dependent methyltransferases; SMART:SM01206:Fibrillarin_2; G3DSA:3.30.200.20:Phosphorylase Kinase, domain 1; G3DSA:3.40.50.150:Vaccinia Virus protein VP39; ProSitePatterns:PS00566:Fibrillarin signature.; PRINTS:PR00052:Fibrillarin signature; MobiDBLite:consensus disorder prediction; Pfam:PF01269:Fibrillarin; GO:0003723:RNA binding; GO:0006364:rRNA processing; GO:0008168:methyltransferase activity; MapolyID:Mapoly0026s0084 | AT4G25630 | ATFIB2,FIB2 | YES |
| Mp2g00370 | -0.6326256 | -0.6843453 | KEGG:K14565:NOP58, nucleolar protein 58; KOG:KOG2572:Ribosome biogenesis protein - Nop58p/Nop5p, [AJ]; G3DSA:1.10.150.460; MobiDBLite:consensus disorder prediction; Pfam:PF01798:snoRNA binding domain, fibrillarin; G3DSA:1.10.246.90; Pfam:PF08156:NOP5NT (NUC127) domain; PTHR10894:SF13; ProSiteProfiles:PS51358:Nop domain profile.; G3DSA:1.10.287.660:Helix hairpin bin; PANTHER:PTHR10894:NUCLEOLAR PROTEIN 5 NUCLEOLAR PROTEIN NOP5 NOP58; Coils:Coil; SUPERFAMILY:SSF89124:Nop domain; SMART:SM00931:NOSIC_2; MapolyID:Mapoly0028s0114 | AT5G27120 | nosymbolavailable | YES |
| Mp4g12650 | -0.6255105 | -0.6759766 | KEGG:K11131:DKC1, NOLA4, CBF5, H/ACA ribonucleoprotein complex subunit 4 [EC:5.4.99.-]; KOG:KOG2529:Pseudouridine synthase, [J]; ProSiteProfiles:PS50890:PUA domain profile.; MobiDBLite:consensus disorder prediction; Pfam:PF16198:tRNA pseudouridylate synthase B C-terminal domain; TIGRFAM:TIGR00451:unchar_dom_2: uncharacterized domain 2; G3DSA:3.30.2350.10:Pseudouridine synthase; Pfam:PF01509:TruB family pseudouridylate synthase (N terminal domain); PTHR23127:SF0:H/ACA RIBONUCLEOPROTEIN COMPLEX SUBUNIT DKC1; SUPERFAMILY:SSF88697:PUA domain-like; G3DSA:2.30.130.70; TIGRFAM:TIGR00425:CBF5: putative rRNA pseudouridine synthase; SMART:SM01136:DKCLD_2; Pfam:PF01472:PUA domain; PANTHER:PTHR23127:CENTROMERE/MICROTUBULE BINDING PROTEIN CBF5; SUPERFAMILY:SSF55120:Pseudouridine synthase; SMART:SM00359:pua_5; Pfam:PF08068:DKCLD (NUC011) domain; CDD:cd02572:PseudoU_synth_hDyskerin; Coils:Coil; GO:0009982:pseudouridine synthase activity; GO:0001522:pseudouridine synthesis; GO:0003723:RNA binding; GO:0006396:RNA processing; GO:0009451:RNA modification; MapolyID:Mapoly0138s0004; | AT3G57150 | NAP57,AtCBF5,CBF5,At NAP57 | YES |
| Mp1g06160 | -0.6134383 | -0.6187978 | KEGG:K14550:UTP10, HEATR1, U3 small nucleolar RNA-associated protein 10; KOG:KOG1837:Uncharacterized conserved protein, C-term missing, [S]; PTHR13457:SF1:HEAT REPEAT-CONTAINING PROTEIN 1; SUPERFAMILY:SSF48371:ARM repeat; PANTHER:PTHR13457:BAP28; Pfam:PF12397:U3 small nucleolar RNA-associated protein 10; SMART:SM01036:BP28CT_2; Pfam:PF08146:BP28CT (NUC211) domain; MapolyID:Mapoly0043s0008 | AT3G06530 | nosymbolavailable | YES |
| Mp1g04040 | -0.6101081 | -0.5781377 | KOG:KOG3070:Predicted RNA-binding protein containing PIN domain and invovled in translation or RNA processing, N-term missing, C-term missing, [J]; CDD:cd04458:CSP_CDS; Pfam:PF00098:Zinc knuckle; ProSitePatterns:PS00352:Cold-shock (CSD) domain signature.; Pfam:PF00313:'Cold-shock' DNA-binding domain; ProSiteProfiles:PS50158:Zinc finger CCHC-type profile.; SMART:SM00343:c2hcfinal6; SUPERFAMILY:SSF57756:Retrovirus zinc finger-like domains; PANTHER:PTHR46565:COLD SHOCK DOMAIN PROTEIN 2; SUPERFAMILY:SSF50249:Nucleic acid-binding proteins; PRINTS:PR00050:Cold shock protein signature; G3DSA:2.40.50.140; G3DSA:4.10.60.10; SMART:SM00357:csp_8; ProSiteProfiles:PS51857:Cold-shock (CSD) domain profile.; GO:0003676:nucleic acid binding; GO:0008270:zinc ion binding; MapolyID:Mapoly0005s0203; | AT4G36020 | CSDP1,AtCSP1,CSP1 | YES |
| Mp7g08650 | -0.6084483 | -0.9226657 | KEGG:K00026:MDH2, malate dehydrogenase [EC:1.1.1.37]; KOG:KOG1494:NAD-dependent malate dehydrogenase, [C]; Pfam:PF02866:lactate/malate dehydrogenase, alpha/beta C-terminal domain; Pfam:PF00056:lactate/malate dehydrogenase, NAD binding domain; ProSitePatterns:PS00068:Malate dehydrogenase active site signature.; G3DSA:3.90.110.10; TIGRFAM:TIGR01772:MDH_euk_gproteo: malate dehydrogenase, NAD-dependent; SUPERFAMILY:SSF51735:NAD(P)-binding Rossmann-fold domains; G3DSA:3.40.50.720; SUPERFAMILY:SSF56327:LDH C-terminal domain-like; CDD:cd01337:MDH_glyoxysomal_mitochondrial; PTHR11540:SF46:MALATE DEHYDROGENASE, CHLOROPLASTIC-RELATED; PANTHER:PTHR11540:MALATE AND LACTATE DEHYDROGENASE; GO:0006099:tricarboxylic acid cycle; GO:0016616:oxidoreductase activity, acting on the CH-OH group of donors, NAD or NADP as acceptor; GO:0030060:L-malate dehydrogenase activity; GO:0003824:catalytic activity; GO:0016491:oxidoreductase activity; GO:0005975:carbohydrate metabolic process; GO:0006108:malate metabolic process; GO:0016615:malate dehydrogenase activity; MapolyID:Mapoly0068s0019 | AT3G47520 | MDH,pNAD-MDH | YES |
| Mp2g07300 | -0.6075863 | -0.7123116 | KEGG:K14815:MRT4, mRNA turnover protein 4; KOG:KOG0816:Protein involved in mRNA turnover, [A]; SUPERFAMILY:SSF160369:Ribosomal protein L10-like; PANTHER:PTHR45841:MRNA TURNOVER PROTEIN 4 MRTO4; Pfam:PF17777:Insertion domain in 60S ribosomal protein L10P; PTHR45841:SF1:MRNA TURNOVER PROTEIN 4 HOMOLOG; CDD:cd05796:Ribosomal_P0_like; G3DSA:3.90.105.20; Pfam:PF00466:Ribosomal protein L10; G3DSA:3.30.70.1730; GO:0042254:ribosome biogenesis; GO:0000027:ribosomal large subunit assembly; MapolyID:Mapoly0015s0017 | AT1G25260 | nosymbolavailable | YES |
| Mp4g09850 | -0.6047301 | -0.6403582 | KEGG:K03111:ssb, single-strand DNA-binding protein; KOG:KOG1653:Single-stranded DNA-binding protein, [L]; CDD:cd04496:SSB_OBF; G3DSA:2.40.50.140; Pfam:PF00436:Single-strand binding protein family; ProSiteProfiles:PS50935:Single-strand binding (SSB) domain profile.; MobiDBLite:consensus disorder prediction; TIGRFAM:TIGR00621:ssb: single-stranded DNA-binding protein; PTHR10302:SF16:NUCLEIC ACID-BINDING, OB-FOLD-LIKE PROTEIN; PANTHER:PTHR10302:SINGLE-STRANDED DNA-BINDING PROTEIN; SUPERFAMILY:SSF50249:Nucleic acid-binding proteins; GO:0006260:DNA replication; GO:0003697:single-stranded DNA binding; MapolyID:Mapoly0132s0028 | AT3G18580 | nosymbolavailable | YES |
| Mp5g07000 | -0.5945964 | -0.6073996 | KEGG:K02331:POL5, MYBBP1A, DNA polymerase phi [EC:2.7.7.7]; KOG:KOG1926:Predicted regulator of rRNA gene transcription (MYB-binding protein), C-term missing, [K]; MobiDBLite:consensus disorder prediction; Pfam:PF04931:DNA polymerase phi; Coils:Coil; SUPERFAMILY:SSF48371:ARM repeat; PANTHER:PTHR13213:MYB-BINDING PROTEIN 1A FAMILY MEMBER; GO:0008134:transcription factor binding; GO:0005730:nucleolus; GO:0003677:DNA binding; GO:0006355:regulation of transcription, DNA-templated; MapolyID:Mapoly0136s0021 | AT5G64420 | nosymbolavailable | YES |
| Mp5g00060 | -0.5930708 | -0.6883725 | KEGG:K14546:UTP5, WDR43, U3 small nucleolar RNA-associated protein 5; KOG:KOG4547:WD40 repeat-containing protein, [R]; Pfam:PF00400:WD domain, G-beta repeat; ProSiteProfiles:PS50294:Trp-Asp (WD) repeats circular profile.; MobiDBLite:consensus disorder prediction; G3DSA:2.130.10.10; Pfam:PF04003:Dip2/Utp12 Family; ProSiteProfiles:PS50082:Trp-Asp (WD) repeats profile.; PANTHER:PTHR45290:OS03G0300300 PROTEIN; PTHR45290:SF1:OS03G0300300 PROTEIN; SUPERFAMILY:SSF50998:Quinoprotein alcohol dehydrogenase-like; SMART:SM00320:WD40_4; GO:0005515:protein binding; MapolyID:Mapoly0078s0006 | AT5G11240 | NuGWD1 | YES |
| Mp4g14950 | -0.5453133 | -0.6254525 | KEGG:K14799:TSR1, pre-rRNA-processing protein TSR1; KOG:KOG1980:Uncharacterized conserved protein, [S]; Pfam:PF08142:AARP2CN (NUC121) domain; SMART:SM01362:DUF663_2; PANTHER:PTHR12858:RIBOSOME BIOGENESIS PROTEIN; MobiDBLite:consensus disorder prediction; ProSiteProfiles:PS51714:Bms1-type guanine nucleotide-binding (G) domain profile.; PTHR12858:SF1:PRE-RRNA-PROCESSING PROTEIN TSR1 HOMOLOG; SMART:SM00785:aarp2cn2; Pfam:PF04950:40S ribosome biogenesis protein Tsr1 and BMS1 C-terminal; GO:0005634:nucleus; GO:0042254:ribosome biogenesis; MapolyID:Mapoly0119s0018 | AT1G42440 | nosymbolavailable | YES |
| Mp6g16880 | -0.5349775 | -0.6461367 | KEGG:K04077:groEL, HSPD1, chaperonin GroEL; KOG:KOG0356:Mitochondrial chaperonin, Cpn60/Hsp60p, [O]; PTHR45633:SF40:CHAPERONIN CPN60-2, MITOCHONDRIAL-LIKE; G3DSA:3.50.7.10:GroEL; CDD:cd03344:GroEL; Pfam:PF00118:TCP-1/cpn60 chaperonin family; Coils:Coil; PRINTS:PR00298:60kDa chaperonin signature; SUPERFAMILY:SSF54849:GroEL-intermediate domain like; SUPERFAMILY:SSF52029:GroEL apical domain-like; ProSitePatterns:PS00296:Chaperonins cpn60 signature.; G3DSA:3.30.260.10:GROEL; Hamap:MF_00600:60 kDa chaperonin [groL].; SUPERFAMILY:SSF48592:GroEL equatorial domain-like; PANTHER:PTHR45633:60 KDA HEAT SHOCK PROTEIN, MITOCHONDRIAL; TIGRFAM:TIGR02348:GroEL: chaperonin GroL; G3DSA:1.10.560.10:GROEL; GO:0006457:protein folding; GO:0016887:ATPase activity; GO:0005524:ATP binding; GO:0042026:protein refolding; MapolyID:Mapoly0144s0025 | AT3G23990 | HSP60,HSP60-3B | YES |

| | | | | | | |
|---|---|---|---|---|---|---|
| Mp4g22190 | -0.5257532 | -0.5273752 | KEGG:K04567:KARS, lysS, lysyl-tRNA synthetase, class II [EC:6.1.1.6]; KOG:KOG1885:Lysyl-tRNA synthetase (class II), [J]; SUPERFAMILY:SSF55681:Class II aaRS and biotin synthetases; SUPERFAMILY:SSF50249:Nucleic acid-binding proteins; MobiDBLite:consensus disorder prediction; PIRSF:PIRSF039101:LysRS2; Pfam:PF01336:OB-fold nucleic acid binding domain; PRINTS:PR00982:Lysyl-tRNA synthetase signature; ProSiteProfiles:PS50862:Aminoacyl-transfer RNA synthetases class-II family profile.; TIGRFAM:TIGR00499:lysS_bact: lysine--tRNA ligase; Coils:Coil; G3DSA:3.30.930.10:Bira Bifunctional Protein, Domain 2; Hamap:MF_00252:Lysine--tRNA ligase [lysS].; PANTHER:PTHR42918:LYSYL-TRNA SYNTHETASE; Pfam:PF00152:tRNA synthetases class II (D, K and N); CDD:cd00775:LysRS_core; CDD:cd04322:LysRS_N; G3DSA:2.40.50.140; GO:0006418:tRNA aminoacylation for protein translation; GO:0003676:nucleic acid binding; GO:0006430:lysyl-tRNA aminoacylation; GO:0004824:lysine-tRNA ligase activity; GO:0000166:nucleotide binding; GO:0005737:cytoplasm; GO:0005524:ATP binding; GO:0004812:aminoacyl-tRNA ligase activity; MapolyID:Mapoly0090s0010 | AT3G11710 | ATKRS-1 | YES |
| Mp2g17870 | -0.5170037 | -0.6987776 | MobiDBLite:consensus disorder prediction; PANTHER:PTHR33356:TIP41-LIKE PROTEIN; PTHR33356:SF5:TIP41-LIKE PROTEIN; MapolyID:Mapoly0094s0056 | AT4G02830 | nosymbolavailable | NO |

Table S2

| Name | Sequence (5'→3') | Usage | Fig |
|---|---|---|---|
| proGC1L-pro-L1 | caccACTTCTGGAGCGAGCGATGAG | Construction of *proGC1L:ELuc* | Fig. 1*B* |
| proGC1L-pro-R2 | ATTGGCCTTATCACAGCAAGGA | | |
| GC1L_tg1-L | ctcgTAAGGCCAATGTGAAGAA | Construction of the genome editing vector for *GC1L* : target 1 | *SI Appendix* , Fig. S3 and S4 |
| GC1L_tg1-R | aaacTTCTTCACATTGGCCTTA | | |
| GC1L_tg2-L | ctcgCATTGACCAGAACGGCAC | Construction of the genome editing vector for *GC1L* : target 2 | *SI Appendix* , Fig. S3 and S4 |
| GC1L_tg2-R | aaacGTGCCGTTCTGGTCAATG | | |
| GC1L-F1 | CCGCGTTAGACGACGAGAT | pRT-PCR | Fig. 1*A* , 6*E* and 6*F* |
| GC1L-R1 | ATTGCAGTCCTCCGCCTTC | | |
| GC1L-F2 | GGATACGGCACCTAGCACAC | pRT-PCR | Fig. 6*C* |
| GC1L-R2 | TGCTCGATCTGCTAACGAACTC | | |
| GCAM1-F1 | GATTGGTGGGCGAATATGGA | pRT-PCR | *SI Appendix* , Fig. S5, S8E and S8F |
| GCAM1-R1 | CATCGGGGTGTAGGAATGGA | | |
| GCAM1-F2 | CCACCTCTCACCACAACACCT | pRT-PCR | Fig. 6*C* |
| GCAM1-R2 | TACCCGGAGTCGCTGAATG | | |
| ERF15-F | AAAGAAGTCTTCCCAAAGTTGC | pRT-PCR | Fig. 6*C* |
| ERF15-R | CTGTTGTTGTTGTTGTTGGTGG | | |
| MpEF1α-F | AAGCCGTCGAAAAGAAGGAG | pRT-PCR | Fig. 1*A* , 6*E* , 6*F* ,<br>*SI Appendix* , S5, S8E and S8F |
| MpEF1α-R | TTCAGGATCGTCCGTTATCC | | |
| MpAPT-F | CGAAAGCCCAAGAAGCTACC | pRT-PCR | Figure 6C |
| MpAPT-R | GTACCCCCGGTTGCAATAAG | | |
